# OCT4 Negatively Regulates the Transcriptional Programming of the Early Region 3 Immune Evasion Genes of Human Adenovirus

**DOI:** 10.1101/2025.04.23.650321

**Authors:** Ashrafali M. Ismail, Amrita Saha, Kimberly A. Morrissey, Dominic Lundquist, Emanuel Garcia, Praveen Balne, Judy L. Cannon, James Chodosh, Jaya Rajaiya

**Author notes:** Corresponding author: Jaya Rajaiya, Ph.D.

## Abstract

Genomes of viruses are constrained by the virions’ nanoscale, and viral nucleotide sequences without function are a luxury. Yet the double-stranded DNA genome of human adenovirus (HAdV) contains large regions without known purpose. Using TRANSFAC and ChIP-Seq analysis, we identified binding of OCT4 (octomer-binding transcription factor 4) to a noncoding region of HAdV-D37 DNA. Manipulation of OCT4 expression impacted viral E3 gene transcription and gp19k protein expression, altering subsequent MHC Class I expression. These effects were specific to OCT4 binding to the adenovirus 5**′** inverted terminal repeat (ITR) within nucleotides 101-159. Using targeted mutations to OCT4, we found one of two OCT4 binding motifs in the ITR to be crucial for repression of E3 gene expression. In OCT4-siRNA treated cells, E3 RID-α gene expression was also upregulated to inhibit pro-apoptotic signals, suggesting that OCT4 binding also indirectly represses viral replication. Consistent with a role for transcription factors in epigenetic modification during infections, OCT4 knockdown also reduced histone H3 acetylation and DNA methylation. In stem cells, OCT4 sustains pluripotency, whereas in somatic cells, OCT4 plays a dispensable role in self-renewal and maintenance. Herein, we show that OCT4 binding also confers a previously unidentified function to non-coding adenovirus DNA.

**Graphical abstract:** 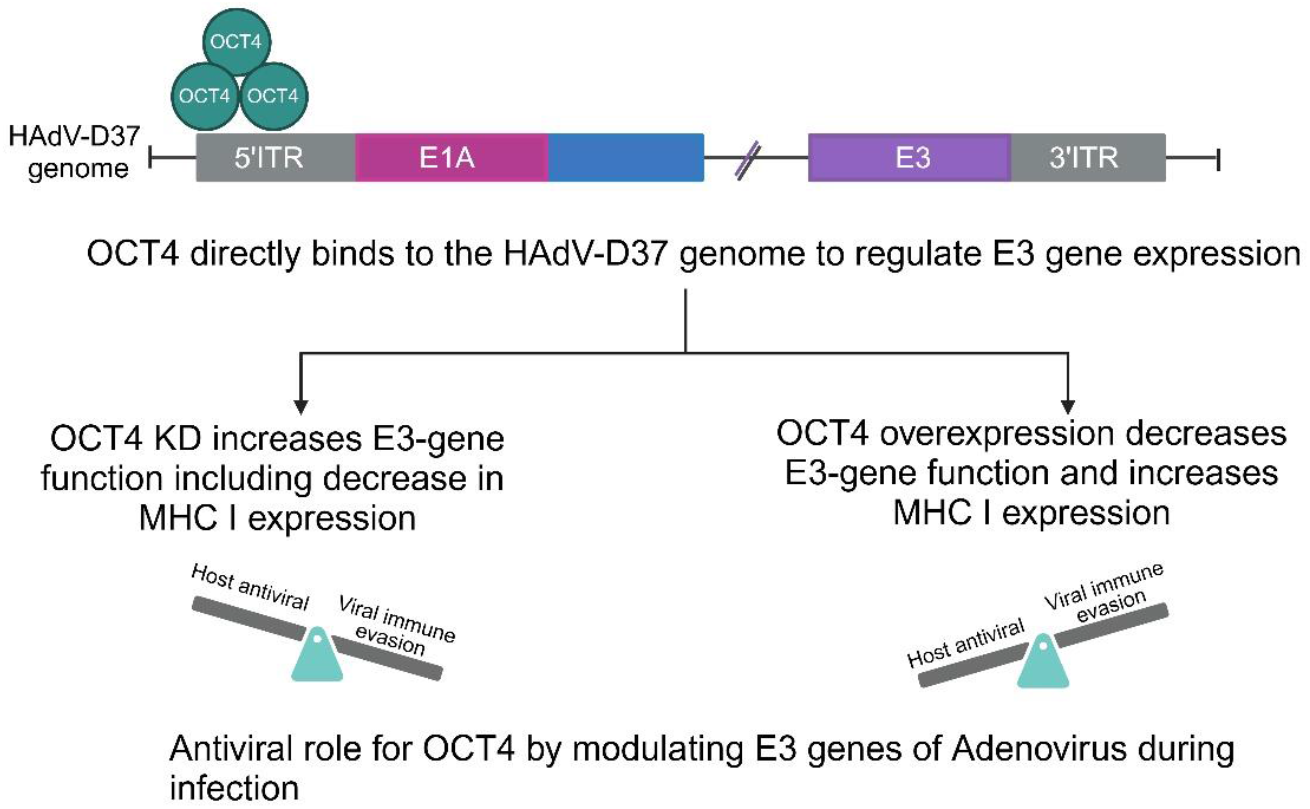

## Introduction

Human adenovirus (HAdV) has been a model organism for scientific discovery, contributing to the landmark findings of p53 inactivation-associated carcinogenesis (1) and RNA splicing (2). The largest and most rapidly evolving HAdV species is HAdV-D (3), with over 70 distinct genotypes, one of which formed the backbone for the Johnson & Johnson COVID-19 vaccine (4). HAdV receptors are expressed on a wide variety of cancers (5), and HAdV has been studied extensively as a chassis for oncolytic virotherapy (6,7). HAdV infection typically results in cell death, potentially amplifying helpful host immune responses to tumor cells (8). Replication of adenovirus DNA occurs in the cell nucleus, and depends on a complex array of viral and host factors (9).

A class V POU protein, OCT4 is a reprogramming transcription factor (TF) indispensable to pluripotency in stem cells (10), with an established role in carcinogenesis (11). In a landmark study, Takahashi and Yamanaka successfully reprogrammed adult fibroblasts into pluripotent stem cells by a combination of transcription factors OCT3/4, SOX2, c-MYC, and KLF4 (12), with c-MYC later shown to be dispensable (13). OCT4 and other members of the POU TF family bind to the octamer motif ATGCAAAT and closely related sequences using a bipartite DNA-binding domain, comprised of two helix-turn-helix domains connected by a variable linker region (14). Although previously thought to be irreversibly inactivated after embryogenesis, OCT4 has been shown to be upregulated in injury, for example in smooth muscle cells of patients with pulmonary hypertension (15), where it appears to be protective (16). OCT4 was was shown to bind to the adenovirus early region 1A (E1A) protein and promote trans-activation of adenoviral gene expression (17). OCT4 has also been shown to play a role in disease associated with other viral infections. For example, OCT4 interaction with the E7 oncogene of human papilloma virus reduces cellular proliferation in cervical cancer cells (18).

HAdV-D37 is a well characterized (19) and highly virulent pathogen, associated with clinically severe infections of the genitourinary tract (20,21) and ocular surface (20). HAdV-D37 has been used extensively in studies of molecular pathogenesis (22-27) and HAdV evolution (28). As part of a larger effort to determine functions for noncoding regions of the HAdV-D37 genome (29), akin to the impetus behind the National Human Genome Research Encyclopedia of DNA Elements (ENCODE) study (30-32), we sought to identify host transcription factor binding sites on HAdD-37 DNA and their potential impact on viral pathogenesis. Herein, we show by *in silico* transcription factor database (TRANSFAC) analysis (33) that OCT4 is highly predicted to bind to the HAdV-D37 genome. Utilizing infection of a pulmonary epithelial cancer cell line, we confirm that OCT4 binds to HAdV-D37 DNA within the inverted terminal repeat (ITR) region to negatively regulate select E3 region gene expression, thus potentially reducing host immune responses to infection, while also limiting viral replication through increased apoptotic cell death. In this work, we also show a direct effect of OCT 4 binding to the ITR on Major Histocompatibility Complex Class I (MHC Class I) expression during infection, in which OCT4 knockdown reduces MHC Class I expression, and OCT4 overexpression increases it. OCT4 binding to the ITR therefore may serve to counteract and balance E1A-driven E3 promoter activation, and potentially provides a novel function for this non-coding region within adenovirus DNA.

## Material and methods

### In silico analysis

Genome-wide binding profiles of TFs in human adenovirus-D37 were predicted using in silico-TRANSFAC (TRANScription FACtor database, version 4.0) analysis (https://genexplain.com/transfac/). TF binding sites were identified on the HadV-D37 genome (GenBank accession no. DQ900900) in comparison to random sequence with the same length and GC content. MEME motif analysis (https://meme-suite.org) was carried out to determine novel OCT4 binding motifs in the HAdV-D37 genome.

### Cells and viruses

A549 cells, a human alveolar carcinoma cell line (ATCC), were cultured in Dulbecco’s modified Eagle medium (DMEM; GIBCO) supplemented with 10% heat-inactivated fetal bovine serum (FBS; GIBCO) and maintained at 37°C with 5% CO2. Human embryonic stem cells (HUES 8) (34), the kind gift of Dr. Dmitry Shvartsman (Harvard Stem Cell Institute), were used as a positive control. HAdV-D37 (ATCC) was grown in A549 cells, purified by CsCl gradient centrifugation, dialyzed, and stored at -80°C. Virus and cell lines were tested repeatedly for endotoxin and mycoplasma contamination prior to use. Virus titration was performed in A549 cells, and the 50% tissue culture infective dose (TCID50) titer was calculated by the formula of Reed and Muench (35)

### ChIP-Seq analysis

A549 cells were infected with HAdV-D37 at 5 moi for 4 hrs and ChIP-Seq was carried out using the iDeal ChIP-seq kit for transcription factors (Diagenode) as per the manufacturer’s instructions. Briefly, virus and mock-infected cells were cross-linked using 1% methanol-free formaldehyde solution (Thermo Fisher Scientific) for 8 min at RT. Excess formaldehyde was quenched by addition of glycine. The cells were washed twice with cold-PBS and collected by scraping in PBS containing protease inhibitor cocktail, and centrifuged at 1600 rpm for 5 min, suspended in sonication buffer, and the chromatin extracts containing cross-linked DNA sonicated (Q700 sonicator, QSonica) to shear DNA to an average fragment size of 100-500 bp. The sheared cross-linked DNA was immunoprecipitated using an ENCODE validated antibody, anti-OCT4, 2 μg (Santa Cruz Biotechnologies, # sc-9081) for ON incubation at 4°C. Negative and positive ChIP controls were performed with rabbit IgG and CTCF antibodies (Diagenode), respectively. ChIP enriched DNA was de-crosslinked, purified using IPure magnetic purification, and relative DNA enrichment verified by quantitative-PCR using control primers targeting human CTCF (positive control) and myoglobin exon (negative control). ChIP-Seq library construction was performed using the illumina TruSeq ChIP sample preparation (Illumina) and AMPure XP beads purification (Beckman Coulter). DNA size selection was performed on a 2% agarose gel and DNA library fragments in the range 175-225 bp were excised, gel purified, and PCR amplified to selectively enrich adapters containing DNA fragments. DNA quality was assessed on the Bioanalyzer High sensitivity Chip, and a sample library prepared for next-generation sequencing (HiSeq). ChIP-Seq data was analyzed using CLC Genomics Workbench (CLCbio, version 7.5). The human reads were aligned using the hg19 reference genome and virus reads aligned with HAdV-D37 reference genome. Each ChiP-Seq experiment was performed in two biological replicates and ChIP-seq peaks with significant more reads than input controls were identified by peak calling using MACS2 (36).

### Electrophoretic mobility shift assay

Nuclear extracts were prepared from A549 cells using NE-PER nuclear and cytoplasmic extraction reagents (Thermo Fisher Scientific). The nuclear extracts were then dialyzed for overnight at 4°C, protein concentrations were determined by BCA protein assay (Thermo Fisher Scientific) and stored at -80°C. Biotin-labeled double-stranded DNA oligos of prototypic OCT4 and its mutants were synthesized by IDT and used for the electrophoretic mobility shift assay (EMSA). Binding reactions were performed using LightShift Chemiluminescent EMSA kit (Thermo Fisher Scientific). Briefly, the reaction mixture contained 10X binding buffer, 50% glycerol, 100mM MgCl2, 1μg/μl Poly (dI.dC) nonspecific competitor DNA, 1%NP-40, 10μg nuclear extract, and 20fmol 38-40 bp DNA probe in a 20μl reaction volume. The mixtures were incubated for 20 min RT. For competition reaction, 200-fold excess of unlabeled target DNA was included prior to the addition of nuclear extract. For supershift assays, 1-4μg of polyclonal antibodies against OCT4 (Abcam) were added at a final concentration of 2 μg/reaction to the binding reaction mixture at the end of the binding reaction and incubated for 20 mins at RT. Samples were separated on 5% native polyacrylamide gels as previously described (37, 38). The electrophoretic reactions were transferred to nylon membranes (Biorad) and cross linked at 120mJ/cm^2^ for 1 min using a UV-light crosslinker (UVP, LLC, Upland CA). Biotin-labeled DNA was detected by chemiluminescence.

### Stable OCT4 expressing cells

pPyCZGIP_OCT4-FLAG-CAG plasmid and empty vector pPyCZGIP_FLAG-CAG were kind gifts from Dr. Yi-Ching Wang (National Cheng Kung University). To generate stable OCT4 and empty vector stable cell lines, pPyCZGIP_OCT4-FLAG-CAG and pPyCZGIP_FLAG-CAG plasmids were transfected into A549 cells using the Lipofectamine 3000 following the manufacturer’s specifications (Thermo Fisher Scientific). According to the observed viability, the cells were allowed to recover for 7-10 days without antibiotics. Selection was then started by adding 0.5 μg/mL puromycin (Thermo Fischer Scientific). Once stable expression of the polyclonal population was verified by OCT4 and FLAG-specific western blot, single-cell clone expansion was initiated with standard growth medium containing 0.5μg/ml puromycin. For limiting dilution, cells were transferred to 96-well suspension plates with ∼1 cell per well. Clonal cell lines were subsequently scaled up, tested for protein expression, and finally preserved in liquid nitrogen.

### Luciferase reporter assay

The wild type and ITR mutants synthesized from IDT (Fig S1)) were cloned into the luciferase reporter plasmid pGL4.16 [luc2] (Promega), in the KpnI and NheI sites after PCR amplification. Amplification was conducted with 100 ng of template adenoviral DNA with the Q5 high fidelity DNA polymerase (New England Biolabs) with a DNA thermal cycler. All constructs were Sanger sequenced for confirmation. Briefly, A549 and OCT4-A549 stable cells were seeded at a density of 1 × 10^4^cells/well in 96-well plates and transiently transfected with the luciferase reporter plasmid constructs (5’ ITR, 3’ ITR wild type or mutant constructs), and luciferase expression measured at 24 hrs using Dual-Glo Luciferase assay system (Promega). Dual-Glo Luciferase assay reagent was added to control and DNA transfected wells in triplicate, incubated at 15 mins RT and firefly luminescence measured on a Spectra Max i3X (Molecular Devices). Following the addition of Dual-Glo Stop and Glo reagent, the plate was incubated for 15 mins RT and then measured for luminescence. The ratio of firefly to Renilla luminescence was calculated and normalized to the ratio from the empty vector control wells.

### Transient transfection

Silencer Select Non-targeting negative control siRNA (Ambion #4390843), OCT4 siRNA-1 ON-TARGETplus Human POU5F1 siRNA, SMARTpool (Horizon Discovery #L-019591-00-0005), were obtained. A reverse transfection method was used. Briefly A549 cells were trypsin treated and the cell suspension was incubated with the negative siRNA or the OCT4-siRNA complex: 50-100pmol of each siRNA with Lipofectamine RNAiMAX (Thermo Fisher Scientific) in Opti-MEM reduced serum medium (Thermo Fischer Scientific). The transfected cells were allowed to grow for 24 hrs in the antibiotic-free growth medium, and were treated with siRNAs for a second time. The transfected cells were then infected with virus, and Western blots performed within 72 hrs of transfection. Luciferase constructs were confirmed by Sanger sequencing. The confirmed clones were transfected using Lipofectamine 3000 (Thermo Fisher Scientific #L3000015). Briefly, 1 day post cell plating, Opti-MEM I medium, 100ng of the specific luciferase construct, 2 ng Renilla vector (at a ratio of 50:1), and p3000 reagent were mixed, and the complex was added to Lipofectamine 3000 and Opti-MEM I medium, and incubated at room temperature for 15mins. This incubated mixture was then added slowly added to the cells.

### RNA isolation and qRT-PCR

A549 cells were transiently transfected by Lipofectamine RNAiMAX. At 72 hrs post-transfection A549 cells were infected with HAdV-D37 at MOI of 1 for 1–24 hrs, washed with PBS, and lysed in TRIzol (Zymo Research). Total RNA was extracted using the Direct-zol RNA kit (Zymo), and the eluent treated with Turbo DNase (Thermo Fisher Scientific) for 30mins at 37°C to remove DNA contamination. Reverse transcription was performed using 100 ng of RNA, oligo(dT), and Moloney murine leukemia virus (M-MLV) reverse transcriptase (Promega). Quantitative real-time PCR (qRT-PCR) analyses was performed with Fast SYBR green mix (Thermo Fisher Scientific) in the QuantStudio 3 system (Thermo Fischer Scientific) using HAdV-D37 primers for viral gene expression kinetics spanning the whole adenoviral genome and human GAPDH as an internal control (Table 1). Each experimental condition was analyzed in triplicate wells and repeated three times. A no template control and an endogenous control (GAPDH) were measured, and the expression levels calculated by the 2^–ddCt method and compared to the mock and infected controls.

### Apoptosis assay

A549 cells were treated with 50 or 100pmol of each siRNA for 24 hrs, and then infected with HAdV-D37 at MOI of 5 for 4 hrs. Infected cells or treated with 1μM Staurosporine (Abcam, #120056) for 4 hrs to induce apoptosis. Cells were then washed with PBS and lysed in cell lysis buffer (Cell Signaling Technology, #9803) with proteinase inhibitors (#5872). For apoptosis assay by western blot, cocktail of pro/p17-caspase-3, cleaved PARP1, muscle actin was used (Abcam #136812). For TUNEL assay, A549 cells were grown to ∼60–80% confluence on slide chambers (Nunc), treated with OCT4 siRNA, and infected with HAdV-D37 at MOI of 5 for 8 hrs. The cells were then fixed in 4% formaldehyde for 10mins, washed in PBS containing 2% FBS, and permeabilized in a solution containing 0.1% Triton X-100 in 2% bovine serum albumin (BSA, Sigma) for 10mins. The TdT reaction was performed according to the manufacturer’s instruction (Click-iT™ TUNEL Alexa Fluor™ Imaging Assay, Thermo Fisher Scientific) Cells were then washed and mounted using Vectashield mounting medium containing DAPI (Vector Labs), and imaged by confocal microscopy using a 63x (NA 1.3) glycerol immersion objective (Leica TCS SP5).

### Western blot analysis

For nuclear extract preparation from siRNA transfection and EMSA experiments, NE-PER extraction reagent (Thermo Fisher Scientific) was used as per the recommended protocol. For whole cell lysates Cell pellets were lysed in cell lysis buffer (Cell Signaling Technology, #9803) with proteinase inhibitors (#5872), and precleared by centrifugation at 14,000g for 10mins at 4°C. Protein concentrations were determined by the BCA Protein Assay (Bio-Rad, #500-0202). Equal Acetyl-Histone H3 - amounts of protein were loaded into NuPAGE 3–8% Tris-acetate protein gels (Thermo Fischer Scientific), and transferred to nitrocellulose membranes (Bio-Rad). Membranes were blocked with 5% BSA, and the primary antibody was incubated overnight at 4°C. Membranes were washed with TBST three times for 10mins each and incubated with secondary antibody for 1 hr at RT. Membranes were washed again and imaged using ChemiDoc (Bio-Rad) and the SuperSignal West Dura Extended Duration Substrate (Thermo Fisher Scientific, #34075). Antibodies used in the study, anti-OCT4 antibody (Abcam # ab19857), Human Adenovirus C Serotype 5 Early E3 18.5 kDa Glycoprotein Antibody (Abnexxa #abx345767), anti-FLAG antibody (Millipore #F1804). Histone modifications were studied using Cell signaling antibodies, Lys9/Lys14 (#9677), Mono-Methyl-Histone H3-Lys4 (#5326), and Di/Tri-Methyl-Histone H3-Lys9 (#5327). HAdV-D E49K antibody was a kind gift from Dr. Hans-Gerhard Burgert, Ludwig-Maximilians-University, Munich 81377, Germany.

### Flow cytometry

A549 cells were washed with PBS without calcium chloride and magnesium chloride (GIBCO, # 70011-044). Cells were detached from culture plates using PBS containing 1 mM EDTA (GIBCO, #15575) to preserve cell-surface HLA proteins. The cells were then incubated with PE-conjugated mouse anti-human HLA-A, B, C antibody (BioLegend #311406), and then washed with PBS and stained with LIVE/DEAD™ Fixable Green Dead Cell Stain (Invitrogen, #L23101A) to assess viability. Cells were then fixed with 2% paraformaldehyde (Electron Microscopy Sciences #15710) and collected using an Attune Nxt Flow Cytometer (Thermo Fisher Scientific). Data were analyzed and visualized using FlowJo software (v10.10).

### Statistical analyses

All assays were performed three times with each sample in triplicate and averages graphed, with error bars indicating the standard deviation (SD). A p-value of < 0.05 was considered statistically significant. Analyses were performed using GraphPad Prism v6.0 (GraphPad Software; San Diego, CA), using Anova and t-test. The asterisk indicates statistical significance (*p-value < 0.05, **p-value < 0.01, ***p-value < 0.001).

### Sequence analyses

Reference sequences for one viral genome from each HAdV species (A-G) [94] were obtained from NCBI (https://www.ncbi.nlm.nih.gov/labs/virus/vssi/#/), and aligned using CLUSTALW (https://www.genome.jp/tools-bin/clustalw). The ITRs were compared using BioEdit (https://bioedit.software.informer.com/7.2/, v7.2.5).

## Results and Discussion

### TRANSFAC analysis

To further current understanding of how adenovirus infection is regulated by host cells, and specifically to identify novel functions for noncoding adenoviral DNA, we first screened for TFs that might bind the adenovirus genome using an *in-silico* TRANSFAC analysis. We identified numerous TF binding sites on the HAdV-D37 genome in comparison to randomly generated DNA sequences with the same length and GC content (Fig. 2A). OCT4 ranked as the top transcription factor to bind the adenovirus genome (ratio=12, p=0.0017), followed by OCT1, ETF, CEBPD, and SP1. Other TFs previously shown to bind the adenovirus genome, including YY1, E2F, and NF-1 (39-42), were also identified in the TRANSFAC analysis (Supplementary Table 1), validating our results. Known functions of OCT4 (Fig. 2B) include maintenance of stem cell pluripotency and tumorigenesis (43). OCT4 was previously shown to interact with adenovirus E1A in pluripotent cells (44), although the importance of this during viral infection in differentiated epithelial cells remains uncertain. We sought to identify a specific function of OCT4 binding to the HAdV-D37 genome, and in Fig. 2C, we show the experimental steps employed to identify TF binding sites on the adenovirus genome through ChIP-Seq.

**Figure 2.**
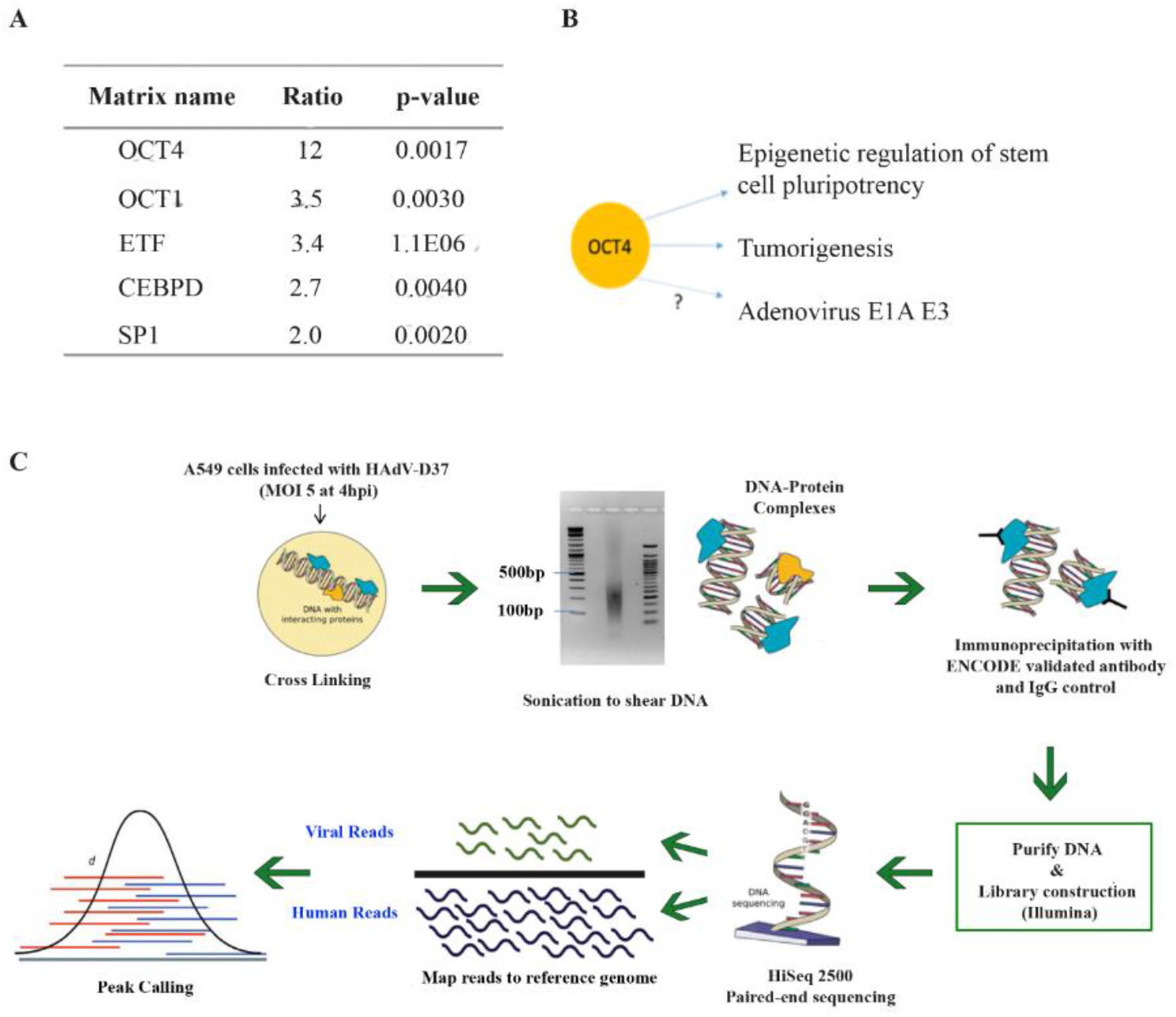
TF binding sites identified on HAdV-D37 genome in comparison to a random DNA sequence with same length and GC content. **(A)** OCT4 binds human adenovirus genome with high affinity. TRANSFAC analysis showing OCT4 as the top transcription factor to bind the adenovirus genome (P=0.0017). **(B)** Known and unknown functions of OCT4. **(C)** ChIP-Seq workflow applied for identification and validation of TF DNA binding sites in the HAdV-D37 genome.

### Hi-Seq analysis of OCT4 targets on the adenovirus genome

To confirm the distribution of OCT4 binding to the adenoviral genome, we performed ChIP-seq on virus infected OCT4 immunoprecipitates, with analyses generated using CLC Genomics Workbench (Qiagen) and Integrated Genomics Viewer, IGV 2.3 (45). Peak calling identified OCT4 binding targets on the following regions of the HAdV-D37 genome (arranged according to nucleotide position): E1A, E1B 55k, E1B-pIX, E2B pol, E2B pTP, L1-52/55kda, L1-pIIIa, L2-pV, L3-hexon, E2A-DBP, L4-100kda, E3-21.8kda CR1α, E3-18.6kda gp19k, E3B-10.4kda RIDα, E3-34kda, and ITR (Fig. 3A and B). Peak calling results were further confirmed by performing fold enrichment assays using real-time PCR. Quantitative PCR performed on ChIP-seq DNA products using primers specific to the adenovirus genes observed in the peak calling confirmed the ChIP-seq results (Fig S1, and Supplemental Table 2). Importantly, no target was identified in the IgG control precipitates.

**Figure 3.**
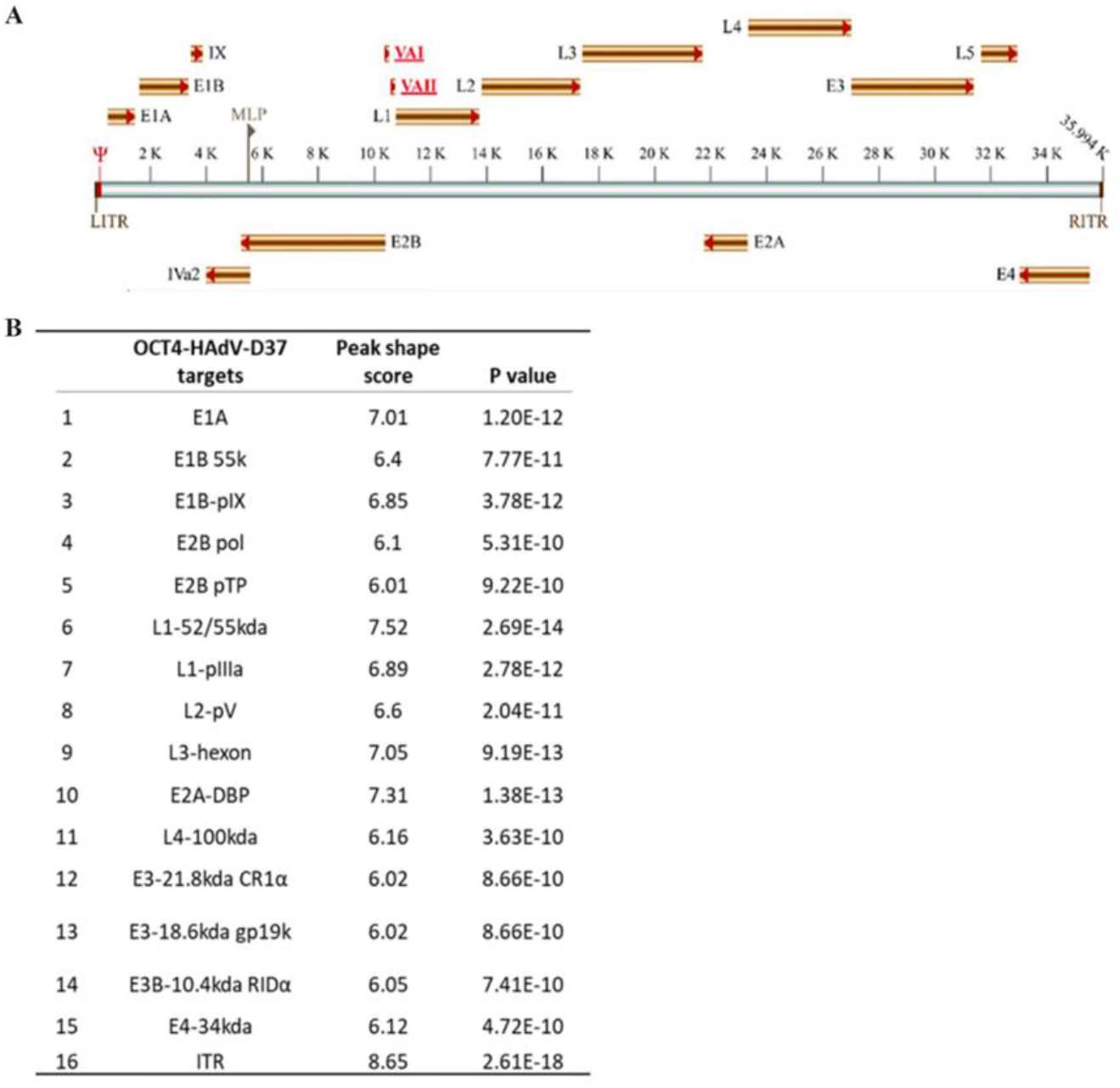
Hi-Seq analysis of OCT4 binding sites on the HAdV-D37 genome **(A)** Transcription map of adenovirus gene targets of OCT4 binding **(B)** Table of OCT4 binding gene targets on the HAdV-D37 genome, identified with using Genomics Viewer, IGV 2.3 (45).

### Specific binding of OCT4 to the ITR

The signature binding motif said to be specific to OCT4 is ATTTGCAT (46,47). Our Chip-Seq and peak calling analysis was supported by the identification of this signature OCT4 binding motif in the 5’-ITR regions (ntd positions 35098-350105), and L2 region (ntd positions 14066-14074). Additionally, the analysis of the peak calling reads discovered other, novel OCT4 binding sites (Fig S2A and S2B) on the adenovirus genome. We validated OCT4 binding to the ITR using EMSA, utilizing a mutation in the specific binding motif that prevented ITR binding by OCT4 (Fig 4). The mutant sequence is shown in the table below.

**Figure 4.**
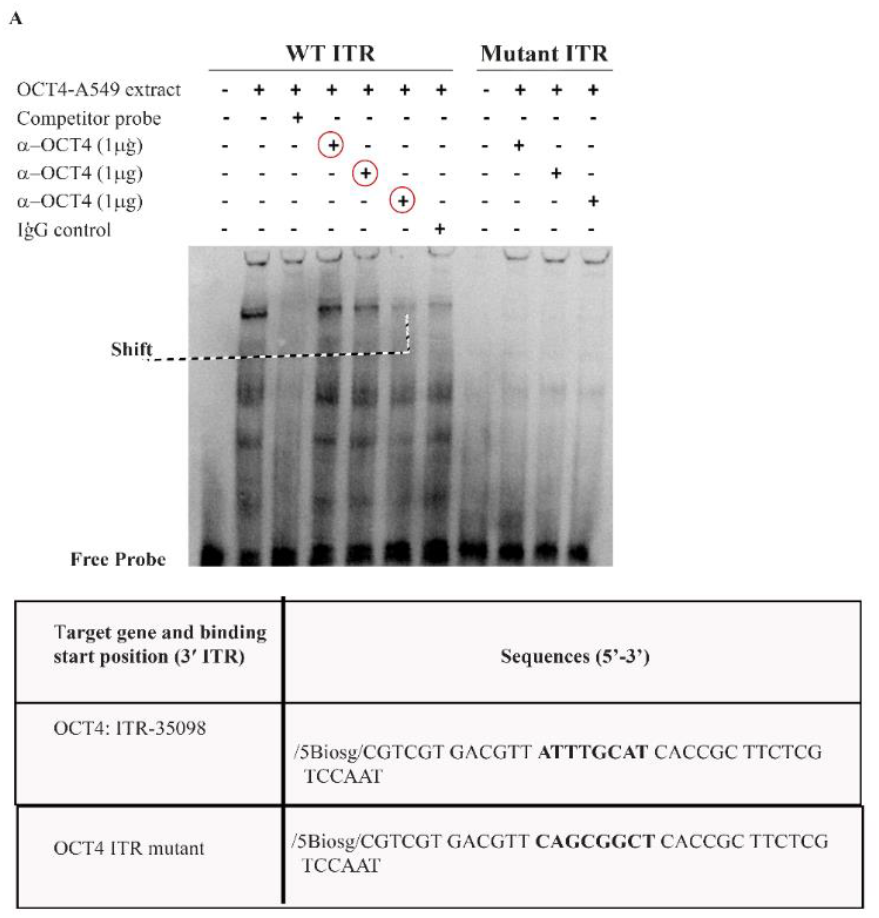
EMSA validation of OCT4 binding to HADV-D37 target “ATTTGCAT”. Mutations made in the binding motif prevented OCT4 binding to ITR (Table)

### OCT4 binding impacts E3 gene expression

We developed several OCT4 siRNAs that suppressed OCT4 mRNA (Fig. 5A) and protein expression (Fig. 5B). Since OCT4 is a multifamily protein, to confirm the specificity of the anti-OCT4 antibody we used an embryonic cell line (HUES 8, Fig. 5B) as a positive control. Further, using quenching techniques by preincubating OCT4 peptide with OCT4 antibody, and then performing Western blots with A549 nuclear extract, we identified a specific OCT4 band (Fig S3). Cells were treated with siRNA for 72 hrs and then infected at an MOI of 5. The expression of individual adenoviral genes was examined by qRT-PCR every hr through 12 hrs post infection using gene specific primers (Supplemental table 3). Significant changes in expression due to OCT4 knockdown were seen in the E3 genes: CR1α, gp19K, and RIDα. Changes were specifically apparent after 4 hrs post infection as compared to the NC-siRNA control. None of the other genes tested showed a statistically significant difference in expression (Fig 5C, S4A). It should be noted that the E3 transcription units in different HAdV species varies in both length and the number of ORFs (28).

**Figure 5.**
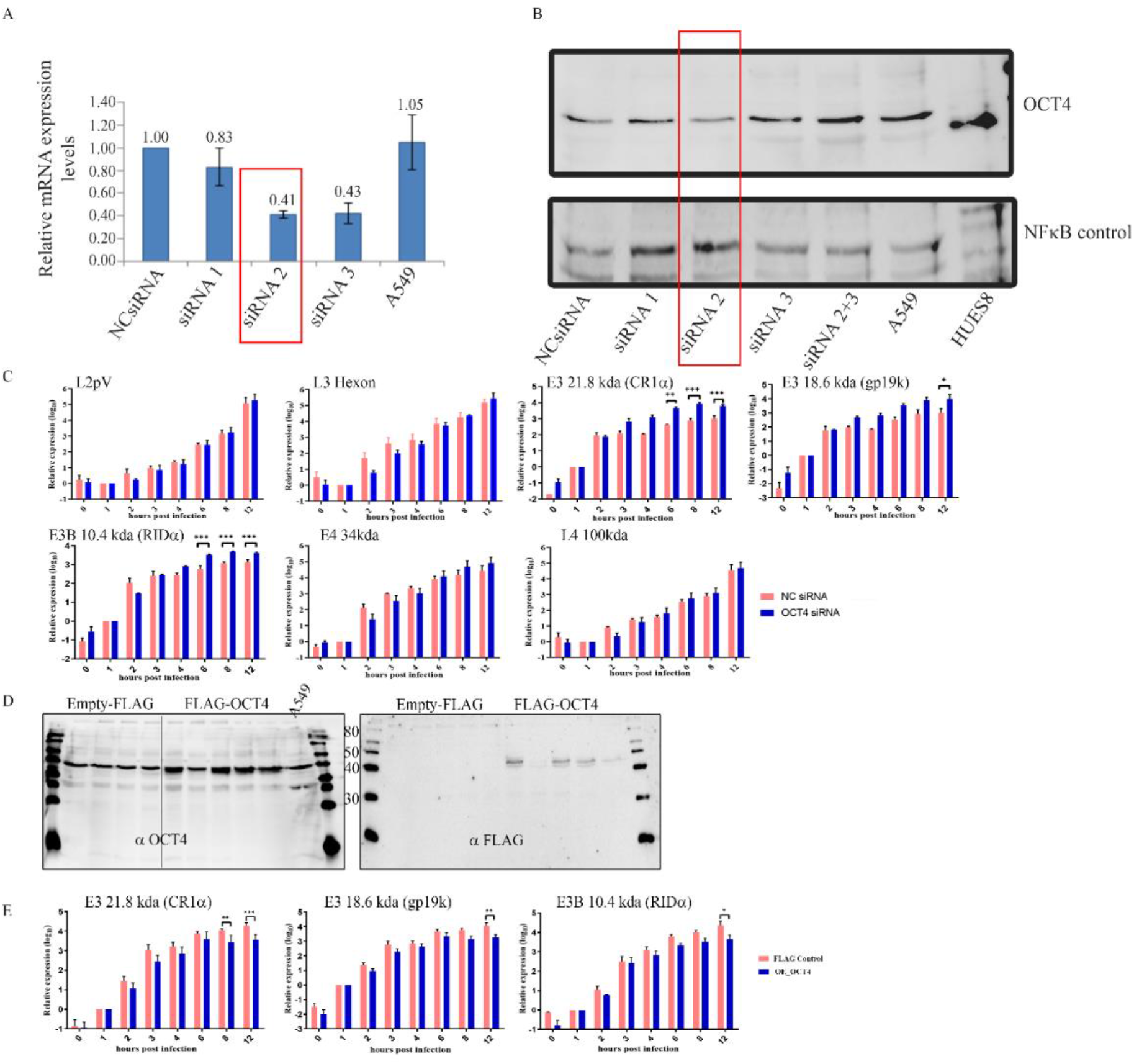
OCT4 regulates immune evasion genes of human adenovirus D37. **(A)** OCT4 mRNA expression levels after 72hrs siRNA treatment. **(B)** Western blot showing OCT4 siRNA knockdown at 72hrs post siRNA treatment. Human embryonic stem cells (HUES8) in which OCT4 is abundantly expressed were used as positive control, and NFkB as a nuclear extract load control. **(C)** Adenovirus gene targets of OCT4 binding and their kinetics of expression at various time points post infection, compared between negative control (NC siRNA) and OCT4-siRNA knockdown in A549 cells, specifically showing changes in expression over time for L2pV, L3 Hexon, CR1α, gp19K, RIDα, E4 34K, and L4100. **(D)** Western blot demonstrating stable expression in cells with constitutive expression of FLAG-OCT4. **(E)** Adenovirus E3 genes CR1α, gp19K, and RIDα expression levels in FLAG and FLAG-OCT4 constitutively expressing cells.

The E3-CR1α protein, previously characterized as 7.1kDa in size for HAdV-C5 and 6.7kDa for HAdV-C2, is a much larger protein in HAdV-D, ∼22kDa (48). The function of CR1α in HAdV-Ds is unconfirmed. However, the 6.7k –CR1α protein from species C was shown to be a transmembrane protein containing an anchor sequence that permits entry into the endoplasmic reticulum, but without a cleavable signal peptide (49). CR1α, together with RIDα and RIDβ cooperate to inhibit TRAIL and thereby reduce the death of infected targeted cells by escaping host cytotoxic effector mechanisms (49). Specifically, 6.7K was shown to affect TRAIL receptor 2 internalization (Lichtenstein DL, et al JVI 2004). E3 gp19K was earlier reported to sequester MHC Class I proteins in the ER, and prevent cytotoxic T cell recognition of HAdV infected cells (50-52). In one study, deletion of gp19k accelerated the clearance of oncolytic adenovirus in a hamster immunocompetent tumor model (53). One of the E3 genes that did not appear in the Chip-seq analysis, and that is uniquely present only in HAdV-D species is the secreted E3-CR1β-49K protein, which binds to leukocytes and suppresses NK cell and T cell function (54). The E3 region in HAdV-D genomes is the most diverse among HAdV species (3,28,48). Our current report showing the increase in CR1α, CR1-gp19K, RIDα, and CR1β gene expression upon OCT4 knockdown suggests that E3 immune evasion functions are regulated by the host transcription factors.

In an OCT 4 expressing stable cell line, OCT4 overexpression specifically reduced the expression of CR1α, gp19K, and RIDα (Fig 5D). Reductions in the expression of other genes was also seen, including L4 100Kda, L1 pIIIa, L3 hexon, and L2pV. (Fig 4E, and Fig S4B). These results strongly suggest that OCT4 negatively regulates specific adenoviral gene expression during natural infection, including some associated with immune evasion by the virus.

### OCT4 indirectly modulates MHC Class I expression during adenovirus infection

Based on the findings above that OCT4 knockdown appeared to increase E3 gene expression and that OCT4 overexpression reduced it, we investigated the effect of OCT4 on MHC Class I expression, a direct measure of immune evasion by adenovirus. It was earlier confirmed that the E3-CR1 gp19k protein sequesters MHC Class I, preventing presentation of antigenic peptides to cytotoxic T lymphocytes (50,55). After 20-24 hrs post infection of OCT4 knockdown cells, MHC Class I expression was reduced compared to NC-siRNA controls in both HAdV-C5 and D37 infection (Fig. 6A). Representative values and MFI histograms are shown in Figure 6B. In stable OCT4 expressing cells however we found increased MHC Class I expression upon HAdV-C5 and D37 infection compared to FLAG control (Fig 6C), also shown in a representative MFI histograms and values (Fig. 6D). These results clearly establish a previously unknown role for OCT4 in regulation of MHC Class I expression in somatic cells, that is further utilized by the infecting pathogens. These experiments utilized a commercial antibody for gp19K that recognizes HAdV-D5 gp19k but is incompletely cross-reactive with the same protein from HAdV-D37. We show increased protein expression of gp19k in siOCT4 cells as compared to NcRNA treated cells after infection (Fig S5A). Similarly, gp19k protein expression was reduced in cells stably transfected to induce OCT4 overexpression (Fig. S5B). E3-CR1β-49K was previously shown to suppress the activation, signaling, and expression of cytokines by T cells, and the activation and cytotoxicity of natural killer cells (54). Using an antibody specific to adenovirus species D 49K, we show that upon OCT4 knockdown, more E49k protein was secreted (Fig. S5C). These data clearly indicate an antiviral role for OCT4 in somatic cells, and a previously unidentified role for the noncoding ITR region of the adenoviral genome.

**Figure 6.**
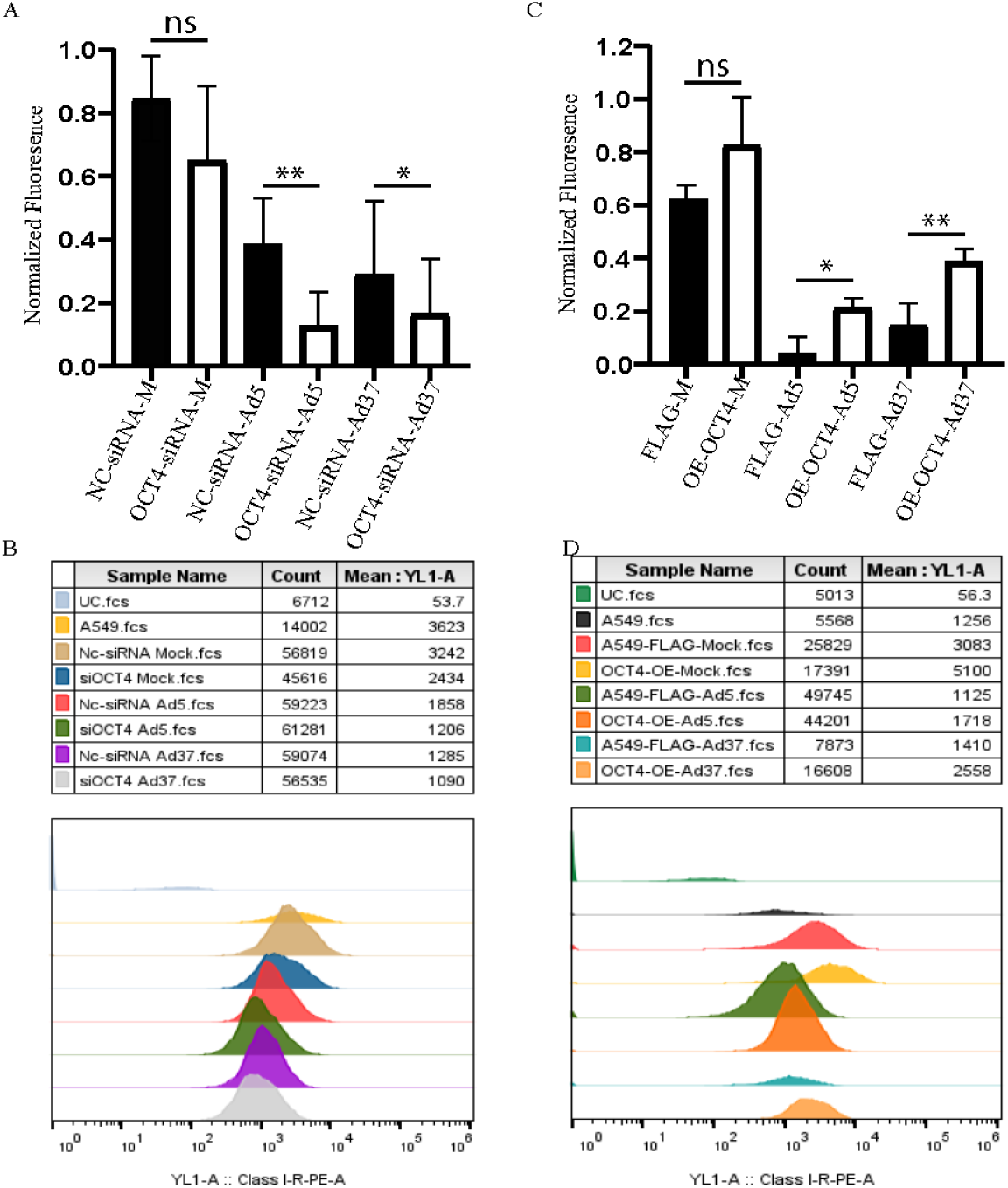
OCT4 regulates E3 gene expression, which in turn modulates MHC-I expression. **(A, B)** OCT4-siRNA treated cells were infected with HAdV-C5 or D37 for 20hrs and stained with an anti-MHC-I antibody and live/dead stain. We gated for live cells and (A) shows the average of mean fluorescence intensity (MFI) of MHC-I from **n=3 (B)** Representative histograms of MHC-I MFI data for (A). **(C, D)** Flag-control (Flag-Ctl) and stable OCT4 overexpression (OE-OCT4) cells were infected with HAdV-C5 or D37 for 20 hrs and stained with an anti-MHC-I antibody and live/dead stain. We gated for live cells and show the MFI of MHC-I **n=3. (D)** Representative histograms of MHC-I MFI data for (C).

### OCT4 binding to the 5’ ITR regulates downstream adenovirus gene expression

In order to understand the specificity of OCT4-ITR binding and its related gene regulation function, mutations were induced in the two regions identified as OCT4 consensus binding sites (ATTTGCAT): positions 43-50 and 110-117. Luciferase assays were conducted on the ITR wild type sequence and its mutants: nucleotide nos. 1-50, 1-50 OCT4-mutant (ATTTGCAT to CAGCGGCT), 101-159, and 101-159 OCT4-binding mutant (ATTTGCAT TO CAGCGGCT), the full length 5’-ITR (wild type), and the full length 5’ITR with both mutations (Fig 7A). Neither the wild type fragment encompassing nucleotides 1-50, and which contains the putative OCT4 binding site within 43-50, nor its mutant activated any luciferase activity. However, the 101-159 fragment showed increased luciferase activity similar to that of the 5’ITR wild type indicating that only 1 of the 2 OCT4 binding sites is needed for transcriptional activity. Furthermore, the indicated mutation in the OCT4 binding site within the 101-159 region reduced luciferase activity similar to that seen with the full length OCT4 binding site mutant (5’ITR mutant) with both mutations (Fig 7B). These results indicate that the OCT4 binding site motif at 43-50 region is dispensable for OCT4-mediated transcriptional activity. To understand potential roles for other transcription factors that may also bind the ITR and regulate adenoviral gene expression, we induced specific point mutations to other predicted transcription factor binding sites (TFBS, Fig. S6A). In the 101-159 fragment, when mutations in binding sites for both OCT4 and ATF-2 were tested, a greater reduction in promoter activity was seen than with the OCT4 binding site mutant alone (Fig. S6B).

**Fig. 7.**
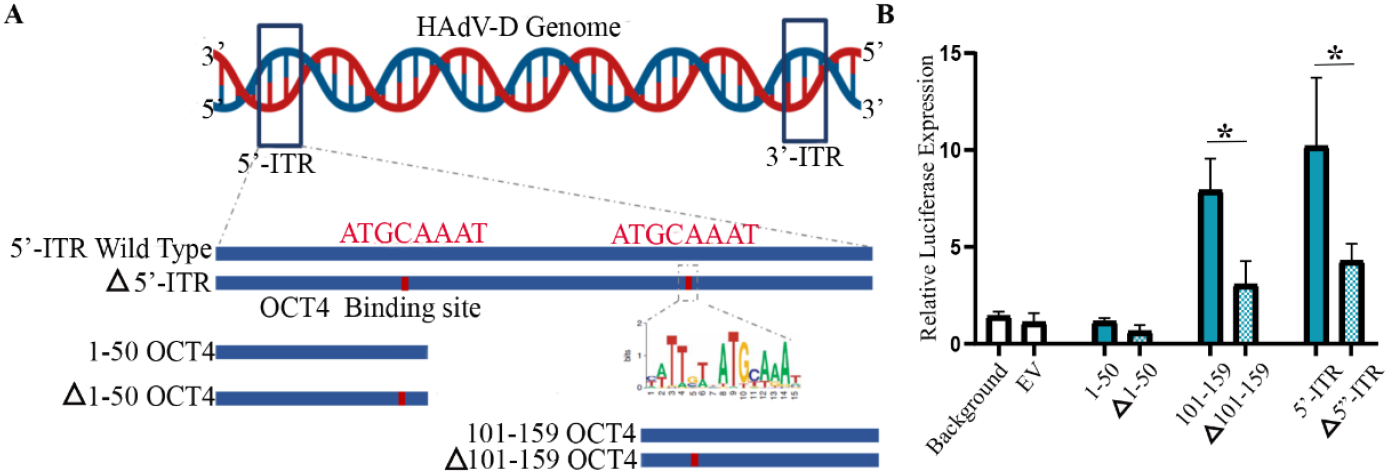
OCT4 ITR site ATTTGCAT at position 110-117 can independently regulate downstream gene expression. **(A)** Schematic of the mutants and OCT4 binding sites **(B)** Luciferase reporter assay showing the potential of OCT4 binding sites within 5′-ITR (1-50, 101-159, and full length) to activate transcription of luciferase gene, and compared with wild type full length. The luciferase activity was predominantly controlled by the 101-159 sequence of the 5’-ITR.

However, in the full length ITR, the OCT4 binding site mutant alone was shown to significantly reduce promoter activity in comparison to the mutations in other TFBS, and with mutation of the OCT4 binding site combined with other TFBS mutations. Cumulatively, these results indicate a premier role for OCT4 in binding of the ITR to regulate adenoviral gene expression. (Mutants sequences are shown in Supplementary Table 4.)

### OCT4 anti-apoptotic activity during adenovirus infection

Another potential outcome of OCT4 binding to the adenovirus genome is its effect on apoptosis, which is tightly regulated during adenovirus infection (56), as with other viruses and non-viral intracellular pathogens (57,58). As above, OCT4 siRNA increased the expression of RIDα potentially reducing apoptosis (59) and indirectly enabling viral replication. To test this idea, we examined the caspase pathway in the setting of OCT4 knockdown followed by infection. OCT4 siRNA pretreated cells showed the presence of cleaved PARP, procaspase 3, and cleaved caspase 3, but not in Nc-siRNA treated controls (Fig 8, left panel). We then treated cells with 1μM Staurosporine to induce apoptosis (60). Cleaved PARP and caspase 3 were both increased and procaspase 3 was considerably reduced in Oct4 siRNA treated cells as compared to NC-siRNA and Staurosporine treated cells (Fig. 8, middle panel). Within 4 hrs of adenovirus infection, OCT4 siRNA treated cells showed reduced caspase activity, absence of cleaved PARP, and a reduced level of cleaved caspase 3 was noted (Fig 8, right panel) as compared to uninfected, OCT4-siRNA transfected (left Vs right panel comparison) or OCT4-siRNA transfected-staurosporin treated cells (middle Vs right panel comparison). In summary, OCT4-siRNA treatment increased the presence of apoptotic markers, while after adenovirus infection apoptotic signals were comparatively reduced, indicating adenoviral gene products counteracted the pro-apoptotic effect of OCT4 knockdown. Specific E3 proteins, most notably RID, downregulate the cell death receptors Fas, TRAIL receptor 1 (TR1) and TR2, and protect infected cells by blocking apoptosis initiated by Fas ligand and TRAIL (49,61-64). As noted above, RIDα expression was increased in OCT4-siRNA treated cells (Fig. 4C), and reduced in OE-OCT4 cells (Fig. 4E). TUNEL assays at 8hrs post infection showed reduced apoptotic signals in OCT4-siRNA treated cells when compared to controls (Fig S7A). In addition, viral replication as determined by titration assay demonstrated increased replication when OCT4 was knocked down (Fig. S7B).

**Figure 8.**
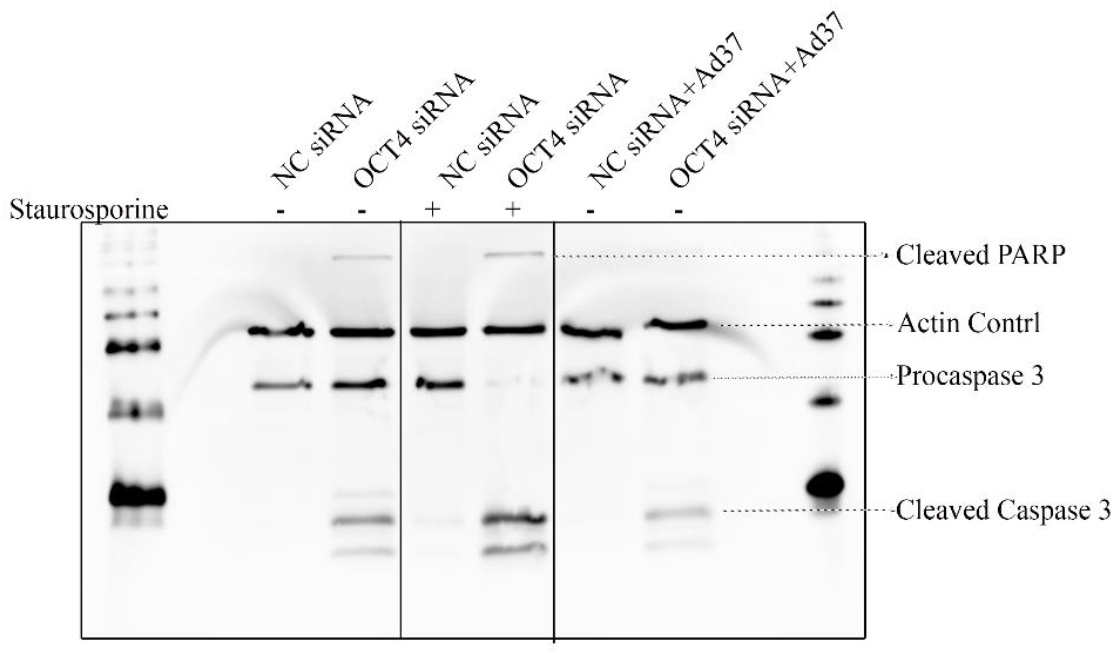
OCT4 knockdown increases apoptosis in the absence of infection, but the impact is reduced in HAdV-D37 infection. OCT4-siRNA treated, uninfected cells (left panel) show increased cleavage of PARP and Caspase 3, indicating initiation of DNA repair and induction of apoptosis. Cleaved PARP and Caspase 3 were further increased in OCT4 knockdown cells treated with staurosporine (middle panel). At 4hrs HAdV-D37 infection (right panel) appeared to compensate for OCT4 knockdown.

### Histone modifications due to adenovirus infection

Histones are highly conserved proteins associated with chromosomal DNA in all eukaryotic cells. They play important roles in the organization and regulation of gene expression, and are major contributors of epigenetic regulation. There are 5 main types of histones divided in two classes, lysine rich-H1, H2A, H2B and arginine rich, H3 and H4. Posttranslational modifications of histones, such as acetylation, methylation, phosphorylation, and ubiquitylation are thought to play a major role in epigenetic regulation. Acetylation of H3-Lys9/14 is primarily associated with transcriptionally active chromatin (65). Methylation of H3-Lys4 is also associated with transcription activation, while tri-methylation of H3-Lys 9 is associated with transcriptional repression (66). Viral proteins can also serve as epigenetic regulators. For example, E1A is known to bind p130 and its associated lysine acetyltransferases, EP300 and CREBBP (67-69) and to induce hypoacetylation (70). In our hands, OCT4-siRNA transfection alone increased acetylation of H3-K9 and 14, methylation of H3-K4me1, and di/tri methylation of H3-K9, but all of these post-translational modification of histones were reduced by adenoviral infection at both 4 and 12hrs post infection (Fig. 9). These latter results clearly indicate that adenovirus infection also regulates epigenetic mechanisms, potentially with long term consequences for cells that can sustain persistent infection with adenoviruses, such as lymphocytes in the tonsils and gastrointestinal tract (71-73).

**Figure 9.**
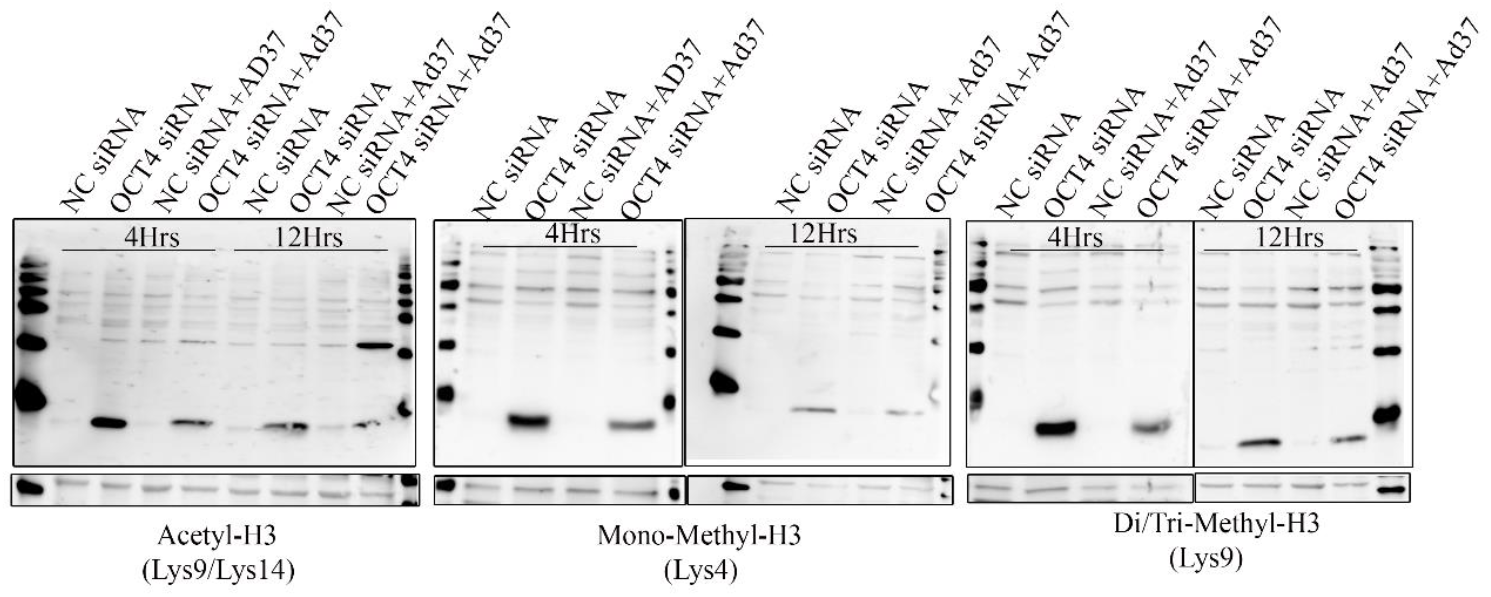
Combinatorial roles for OCT4 and histone modifications. Time-dependent reductions in **(A)** acetylation of H3 (Ly9/14) **(B)** methylation of Mono-Methyl-H3 (Lys4) and **(C)** methylation of Di/Tri-Methyl-H3 (Lys 9) upon OCT4 knockdown followed by HAdV-D37 infection. NFκB was used as for a nuclear control.

**Figure 10.**
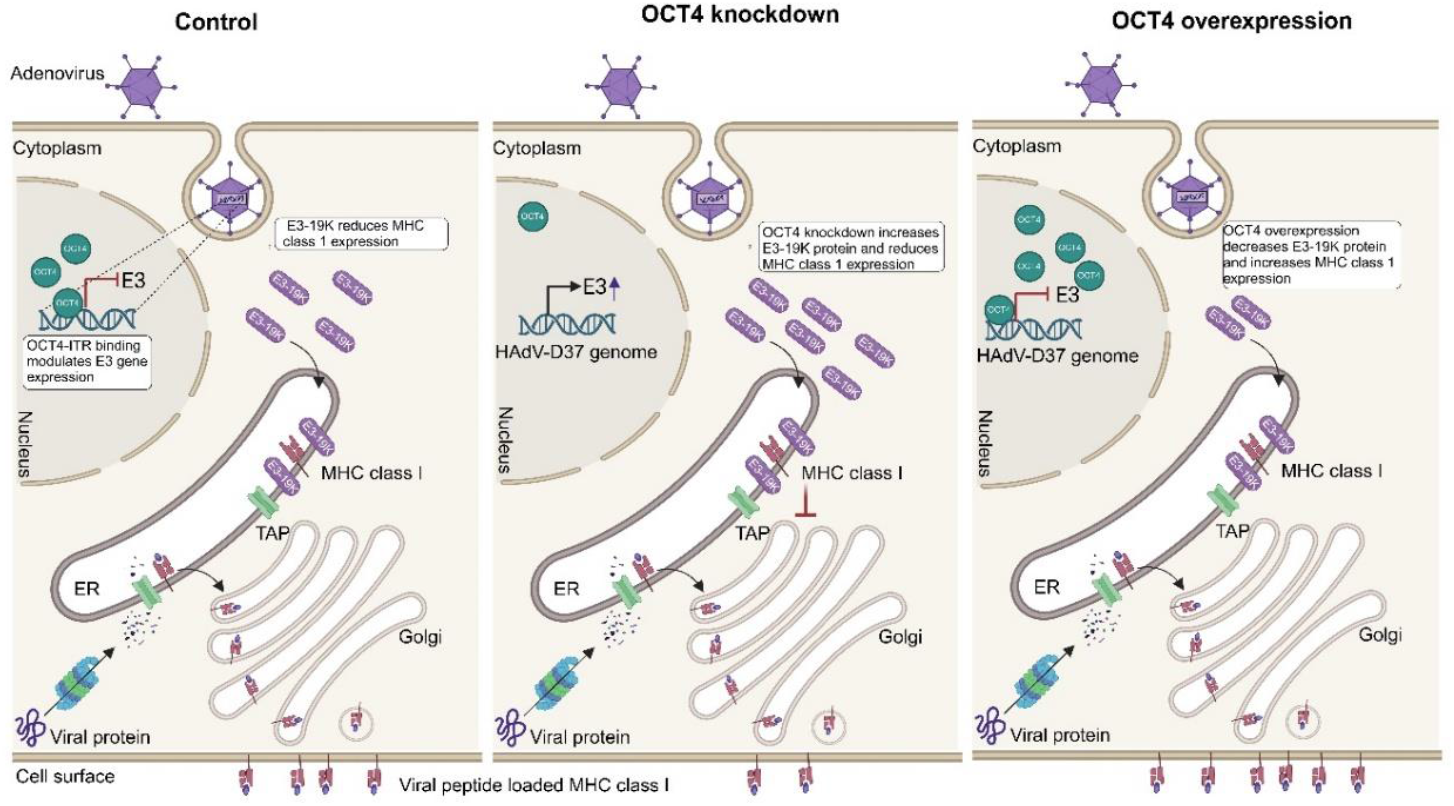
Overall Schematic: The control (left) depicts a normal adenovirus infection. OCT4 knockdown prior to infection (middle panel) increases E3 gene expression, followed by a decrease in MHC Class I expression. OCT4 overexpression (right panel) reduces E3 gene products, and as a result enhances MHC Class I expression.

## Author Contributions

J.R. and A.M.I. conceived the project, designed, conducted experiments and analyzed the data. A.S. supervised and guided for repeat and reproducible data. K.A.M and J.L.C performed and analyzed flow cytometry data. E.G. and D.L prepared sample for flow cytometry and also conducted Western blot experiments. J.C. provided consultation throughout the project, and extensively edited the manuscript. J.R. supervised the project and composed the paper. JR, and JC, provided funding. All authors reviewed the submitted manuscript.

Correspondence and requests for materials should be addressed to Jaya Rajaiya.

## References

1. Levine, A.J. (2009) The common mechanisms of transformation by the small DNA tumor viruses: The inactivation of tumor suppressor gene products: p53. Virology, 384, 285–293.

2. Berk, A.J. and Sharp, P.A. (1978) Structure of the adenovirus 2 early mRNAs. Cell, 14, 695–711.

3. Robinson, C.M., Singh, G., Lee, J.Y., Dehghan, S., Rajaiya, J., Liu, E.B., Yousuf, M.A., Betensky, R.A., Jones, M.S., Dyer, D.W. et al. (2013) Molecular evolution of human adenoviruses. Sci Rep, 3, 1812.

4. Barouch, D.H. (2022) Covid-19 Vaccines - Immunity, Variants, Boosters. N Engl J Med, 387, 1011–1020.

5. Hensen, L.C.M., Hoeben, R.C. and Bots, S.T.F. (2020) Adenovirus Receptor Expression in Cancer and Its Multifaceted Role in Oncolytic Adenovirus Therapy. Int J Mol Sci, 21.

6. Mantwill, K., Klein, F.G., Wang, D., Hindupur, S.V., Ehrenfeld, M., Holm, P.S. and Nawroth, R. (2021) Concepts in Oncolytic Adenovirus Therapy. Int J Mol Sci, 22.

7. Leslie, M. (2023) Tumor-killing viruses score rare success in late-stage trial. Science, 382, 1101–1102.

8. Peter, M. and Kuhnel, F. (2020) Oncolytic Adenovirus in Cancer Immunotherapy. Cancers (Basel),12.

9. Charman, M., Herrmann, C. and Weitzman, M.D. (2019) Viral and cellular interactions during adenovirus DNA replication. FEBS Lett, 593, 3531–3550.

10. Bakhmet, E.I. and Tomilin, A.N. (2021) Key features of the POU transcription factor Oct4 from an evolutionary perspective. Cell Mol Life Sci, 78, 7339–7353.

11. Hochedlinger, K., Yamada, Y., Beard, C. and Jaenisch, R. (2005) Ectopic expression of Oct-4 blocks progenitor-cell differentiation and causes dysplasia in epithelial tissues. Cell, 121, 465–477.

12. Takahashi, K. and Yamanaka, S. (2006) Induction of pluripotent stem cells from mouse embryonic and adult fibroblast cultures by defined factors. Cell, 126, 663–676.

13. Wernig, M., Meissner, A., Cassady, J.P. and Jaenisch, R. (2008) c-Myc is dispensable for direct reprogramming of mouse fibroblasts. Cell Stem Cell, 2, 10–12.

14. Jerabek, S., Merino, F., Scholer, H.R. and Cojocaru, V. (2014) OCT4: dynamic DNA binding pioneers stem cell pluripotency. Biochimica et biophysica acta, 1839, 138–154.

15. Firth, A.L., Yao, W., Remillard, C.V., Ogawa, A. and Yuan, J.X. (2010) Upregulation of Oct-4 isoforms in pulmonary artery smooth muscle cells from patients with pulmonary arterial hypertension. Am J Physiol Lung Cell Mol Physiol, 298, L548–557.

16. Shin, J., Tkachenko, S., Chaklader, M., Pletz, C., Singh, K., Bulut, G.B., Han, Y.M., Mitchell, K., Baylis, R.A., Kuzmin, A.A. et al. (2022) Endothelial OCT4 is atheroprotective by preventing metabolic and phenotypic dysfunction. Cardiovascular research, 118, 2458–2477.

17. Fessler, S.P. and Young, C.S. (1998) Control of adenovirus early gene expression during the late phase of infection. J Virol, 72, 4049–4056.

18. Panayiotou, T., Michael, S., Zaravinos, A., Demirag, E., Achilleos, C. and Strati, K. (2020) Human papillomavirus E7 binds Oct4 and regulates its activity in HPV-associated cervical cancers. PLoS Pathog, 16, e1008468.

19. Robinson, C.M., Shariati, F., Gillaspy, A.F., Dyer, D.W. and Chodosh, J. (2008) Genomic and bioinformatics analysis of human adenovirus type 37: new insights into corneal tropism. BMC genomics, 9, 213.

20. de Jong, J.C., Wigand, R., Wadell, G., Keller, D., Muzerie, C.J., Wermenbol, A.G. and Schaap, G.J. (1981) Adenovirus 37: identification and characterization of a medically important new adenovirus type of subgroup D. J Med Virol, 7, 105–118.

21. Swenson, P.D., Lowens, M.S., Celum, C.L. and Hierholzer, J.C. (1995) Adenovirus types 2, 8, and 37 associated with genital infections in patients attending a sexually transmitted disease clinic. Journal of clinical microbiology, 33, 2728–2731.

22. Chintakuntlawar, A.V., Zhou, X., Rajaiya, J. and Chodosh, J. (2010) Viral capsid is a pathogen-associated molecular pattern in adenovirus keratitis. PLoS Pathog, 6, e1000841.

23. Lee, J.S., Ismail, A.M., Lee, J.Y., Zhou, X., Materne, E.C., Chodosh, J. and Rajaiya, J. (2019) Impact of dynamin 2 on adenovirus nuclear entry. Virology, 529, 43–56.

24. Menon, B.B., Zhou, X., Spurr-Michaud, S., Rajaiya, J., Chodosh, J. and Gipson, I.K. (2016) Epidemic Keratoconjunctivitis-Causing Adenoviruses Induce MUC16 Ectodomain Release To Infect Ocular Surface Epithelial Cells. mSphere, 1.

25. Robinson, C.M., Zhou, X., Rajaiya, J., Yousuf, M.A., Singh, G., DeSerres, J.J., Walsh, M.P., Wong, S., Seto, D., Dyer, D.W. et al. (2013) Predicting the next eye pathogen: analysis of a novel adenovirus. MBio, 4, e00595–00512.

26. Singh, G., Zhou, X., Lee, J.Y., Yousuf, M.A., Ramke, M., Ismail, A.M., Lee, J.S., Robinson, C.M., Seto, D., Dyer, D.W. et al. (2015) Recombination of the epsilon determinant and corneal tropism: Human adenovirus species D types 15, 29, 56, and 69. Virology, 485, 452–459.

27. Walsh, M.P., Chintakuntlawar, A., Robinson, C.M., Madisch, I., Harrach, B., Hudson, N.R., Schnurr, D., Heim, A., Chodosh, J., Seto, D. et al. (2009) Evidence of molecular evolution driven by recombination events influencing tropism in a novel human adenovirus that causes epidemic keratoconjunctivitis. PLoS One, 4, e5635.

28. Robinson, C.M., Seto, D., Jones, M.S., Dyer, D.W. and Chodosh, J. (2011) Molecular evolution of human species D adenoviruses. Infect Genet Evol, 11, 1208–1217.

29. Ismail, A.M., Lee, J.S., Lee, J.Y., Singh, G., Dyer, D.W., Seto, D., Chodosh, J. and Rajaiya, J. (2018) Adenoviromics: Mining the Human Adenovirus Species D Genome. Front Microbiol, 9, 2178.

30. Qu, H. and Fang, X. (2013) A brief review on the Human Encyclopedia of DNA Elements (ENCODE) project. Genomics Proteomics Bioinformatics, 11, 135–141.

31. Consortium, E.P., Birney, E., Stamatoyannopoulos, J.A., Dutta, A., Guigo, R., Gingeras, T.R., Margulies, E.H., Weng, Z., Snyder, M., Dermitzakis, E.T. et al. (2007) Identification and analysis of functional elements in 1% of the human genome by the ENCODE pilot project. Nature, 447, 799–816.

32. M, K.P., Shyama, S.K., D’Costa, A., Kadam, S.B., Sonaye, B.H. and Chaubey, R.C. (2017) Evaluation of DNA damage induced by gamma radiation in gill and muscle tissues of Cyprinus carpio and their relative sensitivity. Ecotoxicol Environ Saf, 144, 166–170.

33. Knuppel, R., Dietze, P., Lehnberg, W., Frech, K. and Wingender, E. (1994) TRANSFAC retrieval program: a network model database of eukaryotic transcription regulating sequences and proteins. J Comput Biol, 1, 191–198.

34. Cowan, C.A., Klimanskaya, I., McMahon, J., Atienza, J., Witmyer, J., Zucker, J.P., Wang, S., Morton, C.C., McMahon, A.P., Powers, D. et al. (2004) Derivation of embryonic stem-cell lines from human blastocysts. N Engl J Med, 350, 1353–1356.

35. Reed, L.J. and Muench, H. (1938) A simple method for estimating fifty per cent endpoints. American journal of epidemiology, 27, 493–497.

36. Zhang, Y., Lin, Z., Xia, Q., Zhang, M. and Zhang, X. (2008) Characteristics and analysis of simple sequence repeats in the cotton genome based on a linkage map constructed from a BC1 population between Gossypium hirsutum and G. barbadense. Genome, 51, 534–546.

37. Rajaiya, J., Hatfield, M., Nixon, J.C., Rawlings, D.J. and Webb, C.F. (2005) Bruton’s tyrosine kinase regulates immunoglobulin promoter activation in association with the transcription factor Bright. Mol Cell Biol, 25, 2073–2084.

38. Rajaiya, J., Nixon, J.C., Ayers, N., Desgranges, Z.P., Roy, A.L. and Webb, C.F. (2006) Induction of immunoglobulin heavy-chain transcription through the transcription factor Bright requires TFII-I. Mol Cell Biol, 26, 4758–4768.

39. Lee, J.S., Galvin, K.M., See, R.H., Eckner, R., Livingston, D., Moran, E. and Shi, Y. (1995) Relief of YY1 transcriptional repression by adenovirus E1A is mediated by E1A-associated protein p300. Genes Dev, 9, 1188–1198.

40. Lee, J.S., See, R.H., Galvin, K.M., Wang, J. and Shi, Y. (1995) Functional interactions between YY1 and adenovirus E1A. Nucleic Acids Res, 23, 925–931.

41. Bruder, J.T. and Hearing, P. (1989) Nuclear factor EF-1A binds to the adenovirus E1A core enhancer element and to other transcriptional control regions. Mol Cell Biol, 9, 5143–5153.

42. Gounari, F., De Francesco, R., Schmitt, J., van der Vliet, P., Cortese, R. and Stunnenberg, H. (1990) Amino-terminal domain of NF1 binds to DNA as a dimer and activates adenovirus DNA replication. EMBO J, 9, 559–566.

43. Huyghe, A., Trajkova, A. and Lavial, F. (2023) Cellular plasticity in reprogramming, rejuvenation and tumorigenesis: a pioneer TF perspective. Trends Cell Biol.

44. Scholer, H.R., Ciesiolka, T. and Gruss, P. (1991) A nexus between Oct-4 and E1A: implications for gene regulation in embryonic stem cells. Cell, 66, 291–304.

45. Robinson, J.T., Thorvaldsdottir, H., Winckler, W., Guttman, M., Lander, E.S., Getz, G. and Mesirov, J.P. (2011) Integrative genomics viewer. Nat Biotechnol, 29, 24–26.

46. Petryniak, B., Staudt, L.M., Postema, C.E., McCormack, W.T. and Thompson, C.B. (1990) Characterization of chicken octamer-binding proteins demonstrates that POU domain-containing homeobox transcription factors have been highly conserved during vertebrate evolution. Proc Natl Acad Sci U S A, 87, 1099–1103.

47. Zhao, F.Q. (2013) Octamer-binding transcription factors: genomics and functions. Front Biosci (Landmark Ed), 18, 1051–1071.

48. Singh, G., Robinson, C.M., Dehghan, S., Jones, M.S., Dyer, D.W., Seto, D. and Chodosh, J. (2013) Homologous recombination in E3 genes of human adenovirus species D. J Virol, 87, 12481–12488.

49. Benedict, C.A., Norris, P.S., Prigozy, T.I., Bodmer, J.L., Mahr, J.A., Garnett, C.T., Martinon, F., Tschopp, J., Gooding, L.R. and Ware, C.F. (2001) Three adenovirus E3 proteins cooperate to evade apoptosis by tumor necrosis factor-related apoptosis-inducing ligand receptor-1 and -2. The Journal of biological chemistry, 276, 3270–3278.

50. Andersson, M., Paabo, S., Nilsson, T. and Peterson, P.A. (1985) Impaired intracellular transport of class I MHC antigens as a possible means for adenoviruses to evade immune surveillance. Cell, 43, 215–222.

51. Burgert, H.G. and Kvist, S. (1985) An adenovirus type 2 glycoprotein blocks cell surface expression of human histocompatibility class I antigens. Cell, 41, 987–997.

52. Rawle, F.C., Tollefson, A.E., Wold, W.S. and Gooding, L.R. (1989) Mouse anti-adenovirus cytotoxic T lymphocytes. Inhibition of lysis by E3 gp19K but not E3 14.7K. J Immunol, 143, 2031–2037.

53. Bortolanza, S., Bunuales, M., Alzuguren, P., Lamas, O., Aldabe, R., Prieto, J. and Hernandez-Alcoceba, R. (2009) Deletion of the E3-6.7K/gp19K region reduces the persistence of wild-type adenovirus in a permissive tumor model in Syrian hamsters. Cancer gene therapy, 16, 703–712.

54. Windheim, M., Southcombe, J.H., Kremmer, E., Chaplin, L., Urlaub, D., Falk, C.S., Claus, M., Mihm, J., Braithwaite, M., Dennehy, K. et al. (2013) A unique secreted adenovirus E3 protein binds to the leukocyte common antigen CD45 and modulates leukocyte functions. Proc Natl Acad Sci U S A, 110, E4884–4893.

55. Paabo, S., Weber, F., Nilsson, T., Schaffner, W. and Peterson, P.A. (1986) Structural and functional dissection of an MHC class I antigen-binding adenovirus glycoprotein. EMBO J, 5, 1921–1927.

56. White, E. (2001) Regulation of the cell cycle and apoptosis by the oncogenes of adenovirus. Oncogene, 20, 7836–7846.

57. Otero-Carrasco, B., Ugarte Carro, E., Prieto-Santamaria, L., Diaz Uzquiano, M., Caraca-Valente Hernandez, J.P. and Rodriguez-Gonzalez, A. (2024) Identifying patterns to uncover the importance of biological pathways on known drug repurposing scenarios. BMC genomics, 25, 43.

58. Clarke, P. and Tyler, K.L. (2009) Apoptosis in animal models of virus-induced disease. Nat Rev Microbiol, 7, 144–155.

59. McNees, A.L., Garnett, C.T. and Gooding, L.R. (2002) The adenovirus E3 RID complex protects some cultured human T and B lymphocytes from Fas-induced apoptosis. J Virol, 76, 9716–9723.

60. Bertrand, R., Solary, E., O’Connor, P., Kohn, K.W. and Pommier, Y. (1994) Induction of a common pathway of apoptosis by staurosporine. Exp Cell Res, 211, 314–321.

61. Elsing, A. and Burgert, H.G. (1998) The adenovirus E3/10.4K-14.5K proteins down-modulate the apoptosis receptor Fas/Apo-1 by inducing its internalization. Proc Natl Acad Sci U S A, 95, 10072–10077.

62. Shisler, J., Yang, C., Walter, B., Ware, C.F. and Gooding, L.R. (1997) The adenovirus E3-10.4K/14.5K complex mediates loss of cell surface Fas (CD95) and resistance to Fas-induced apoptosis. J Virol, 71, 8299–8306.

63. Tollefson, A.E., Hermiston, T.W., Lichtenstein, D.L., Colle, C.F., Tripp, R.A., Dimitrov, T., Toth, K., Wells, C.E., Doherty, P.C. and Wold, W.S. (1998) Forced degradation of Fas inhibits apoptosis in adenovirus-infected cells. Nature, 392, 726–730.

64. Tollefson, A.E., Toth, K., Doronin, K., Kuppuswamy, M., Doronina, O.A., Lichtenstein, D.L., Hermiston, T.W., Smith, C.A. and Wold, W.S. (2001) Inhibition of TRAIL-induced apoptosis and forced internalization of TRAIL receptor 1 by adenovirus proteins. J Virol, 75, 8875–8887.

65. Balakrishnan, L. and Milavetz, B. (2007) Histone hyperacetylation in the coding region of chromatin undergoing transcription in SV40 minichromosomes is a dynamic process regulated directly by the presence of RNA polymerase II. J Mol Biol, 365, 18–30.

66. Black, J.C., Van Rechem, C. and Whetstine, J.R. (2012) Histone lysine methylation dynamics: establishment, regulation, and biological impact. Mol Cell, 48, 491–507.

67. Sherr, C.J. and McCormick, F. (2002) The RB and p53 pathways in cancer. Cancer Cell, 2, 103–112.

68. Berk, A.J. (2005) Recent lessons in gene expression, cell cycle control, and cell biology from adenovirus. Oncogene, 24, 7673–7685.

69. Ferrari, R., Pellegrini, M., Horwitz, G.A., Xie, W., Berk, A.J. and Kurdistani, S.K. (2008) Epigenetic reprogramming by adenovirus e1a. Science, 321, 1086–1088.

70. Horwitz, G.A., Zhang, K., McBrian, M.A., Grunstein, M., Kurdistani, S.K. and Berk, A.J. (2008) Adenovirus small e1a alters global patterns of histone modification. Science, 321, 1084–1085.

71. Kosulin, K., Geiger, E., Vecsei, A., Huber, W.D., Rauch, M., Brenner, E., Wrba, F., Hammer, K., Innerhofer, A., Potschger, U. et al. (2016) Persistence and reactivation of human adenoviruses in the gastrointestinal tract. Clin Microbiol Infect, 22, 381 e381–381 e388.

72. Zhang, Y., Huang, W., Ornelles, D.A. and Gooding, L.R. (2010) Modeling adenovirus latency in human lymphocyte cell lines. J Virol, 84, 8799–8810.

73. Garnett, C.T., Talekar, G., Mahr, J.A., Huang, W., Zhang, Y., Ornelles, D.A. and Gooding, L.R. (2009) Latent species C adenoviruses in human tonsil tissues. J Virol, 83, 2417–2428.

74. Hatfield, L. and Hearing, P. (1993) The NFIII/OCT-1 binding site stimulates adenovirus DNA replication in vivo and is functionally redundant with adjacent sequences. J Virol, 67, 3931–3939.

75. Zhou, J.J., Meng, Z., Zhou, Y., Cheng, D., Ye, H.L., Zhou, Q.B., Deng, X.G. and Chen, R.F. (2016) Hepatitis C virus core protein regulates OCT4 expression and promotes cell cycle progression in hepatocellular carcinoma. Oncol Rep, 36, 582–588.

