## Supplemental Fig and Table for "OCT4 Negatively Regulates the Transcriptional Programming of the Early Region 3 Immune Evasion Genes of Human Adenovirus"

### Supplemental Figures

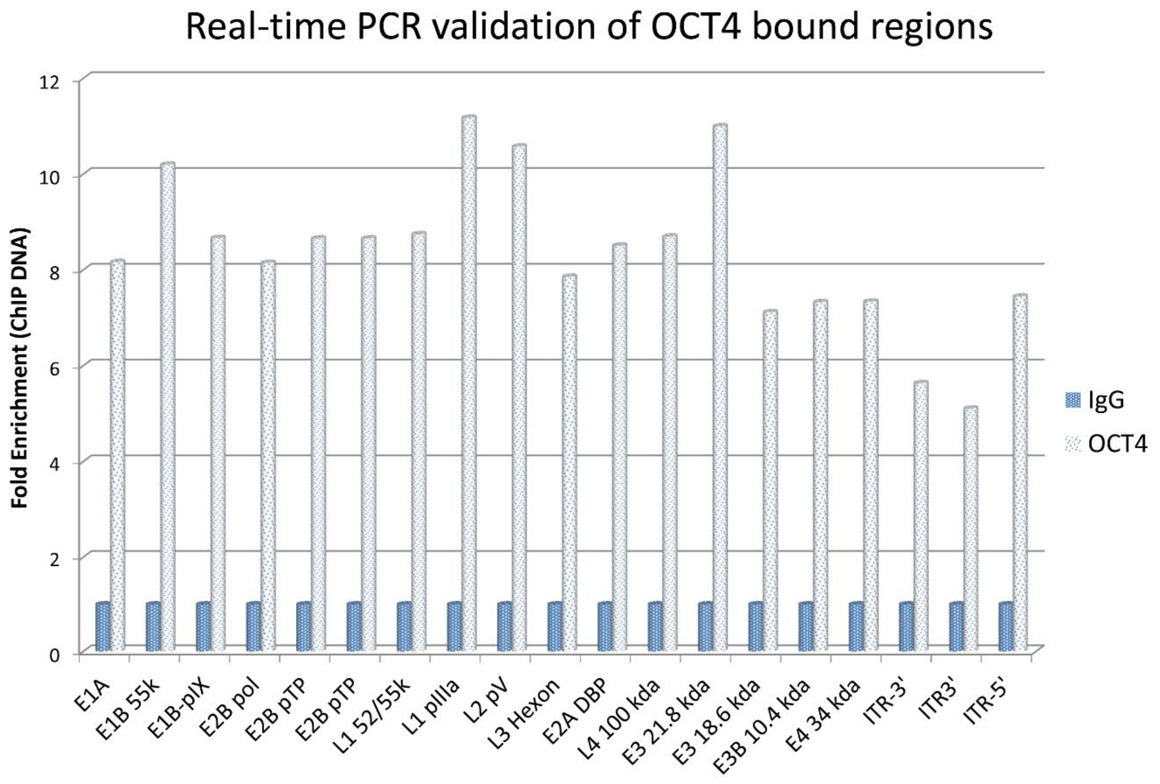

**Figure S1:** Real time PCR using primers specific for adenovirus gene targets indicated in Chip-Seq peak calling analysis. The corresponding enrichment assay for the IgG control showed no target genes were amplified.

A

| HAdV-D37 region | Strand | Start | *p*-value | | Sites | |
|---|---|---|---|---|---|---|
| L2-pV_15898_15998 | - | 38 | 2.22e-4 | CGGGCGTGA | ACACCA | CCGCCGTGCC |
| E2A-DBP_22405_22505 | + | 2 | 2.22e-4 | G | ACACCA | GGGGTCGCGG |
| E2B-pTP_9173_9273 | - | 69 | 5.33e-4 | GCCGTCACCG | AAACCA | TGCGTCGCCG |
| 5'ITR_34564_34664 | + | 54 | 5.33e-4 | AGGTGAAGC A | ATACCA | CCCCATGCGG |
| E1B-pIX_3361_3461 | + | 88 | 6.89e-4 | TTTTCTGCAG | ACATCA | TGAACGG |

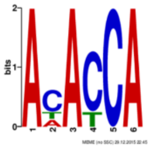

| HAdV-D37 region | Strand | Start | *p*-value | | Sites | |
|---|---|---|---|---|---|---|
| E1B-pIX_3361_3461 | - | 36 | 3.88e-6 | CCGCCTTTAT | ACTCACATT | AGCCACGCCC |
| L2-pV_34564_34664 | + | 34 | 3.88e-6 | GAACCAGCAC | ACTCACTTT | TAGGTGAAGC |

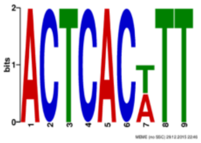

Novel OCT4 binding motifs identified in the HAdV-D37 genome using MEME motif analysis

B

OCT4 peak calling analysis (Expt-1 and -2)

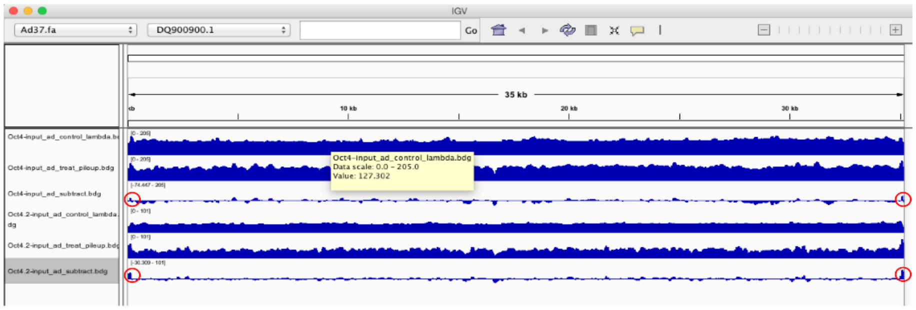

○ The regions marked in red above are the Inverted terminal repeats (ITR) in the virus genome. i.e, they are complementary sequences. This two region also contain the known OCT4 binding motif "ATTTGCAT"

**Figure S2: (A)** Novel OCT4 binding sites on the both strands of the adenovirus genome, their region and motifs. **(B)** Raw peak calling analysis of OCT4 binding to the HAdV-D37 genome.

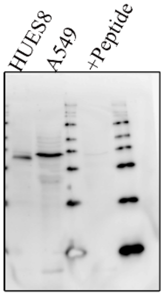

**Figure S3:** To confirm specificity of the OCT4 band seen in A549 cells, the lysate was compared to lysates from embryonic cell line HUES8. In addition, the OCT4 antibody was incubated with OCT4 peptide prior to use to test for the specificity of OCT4 band.

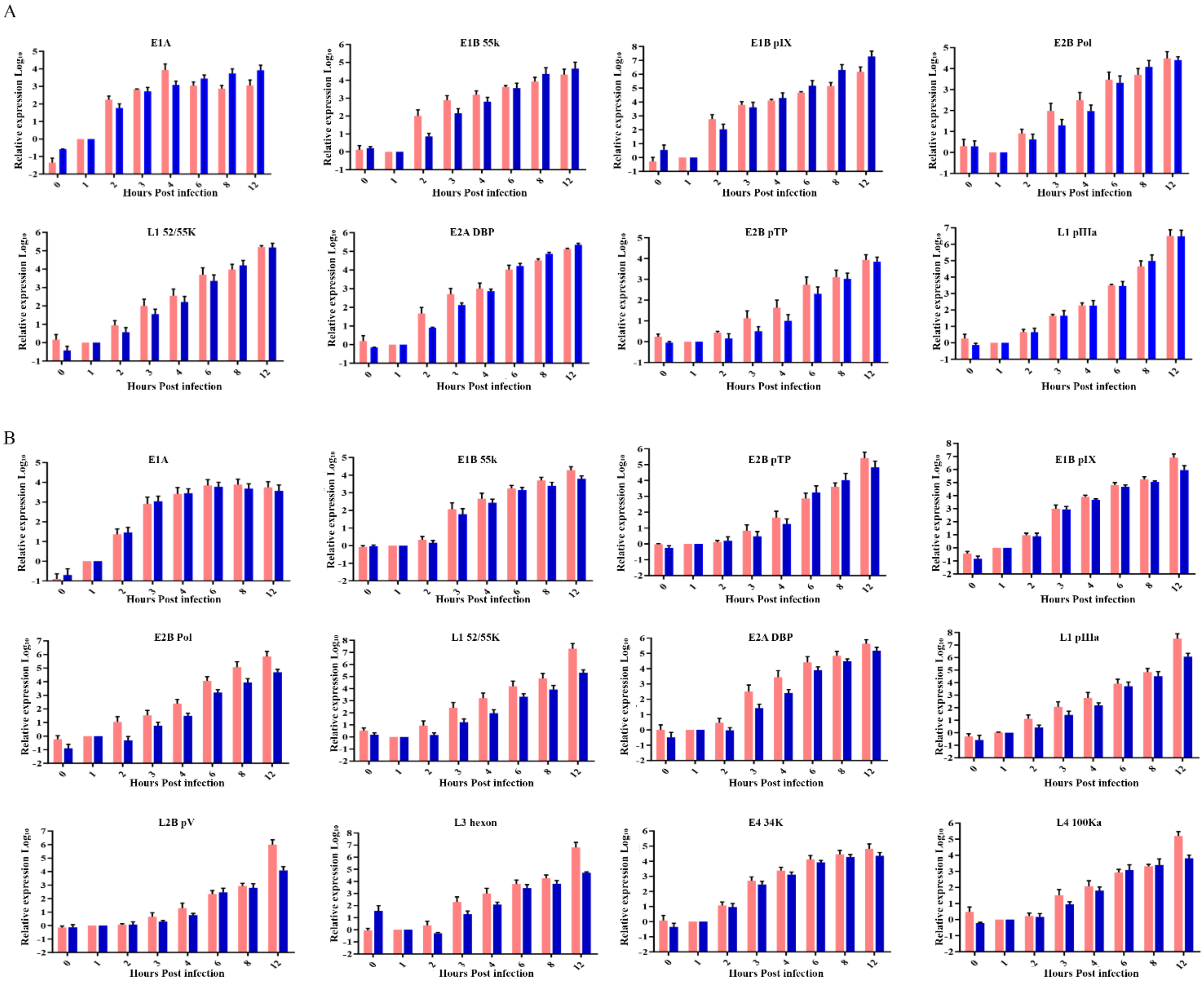

**Figure S4:** OCT4 regulates gene expression from the adenovirus genome. **(A)** OCT4-siRNA treated cells infected with HAdV-D37 were compared at various time points by qRT-PCR for the adenoviral gene transcripts identified by peak calling analysis. **(B)** Cells stably expressing OCT4 were infected with HAdV-D37, and adenovirus gene transcripts measured at the indicated time points post infection.

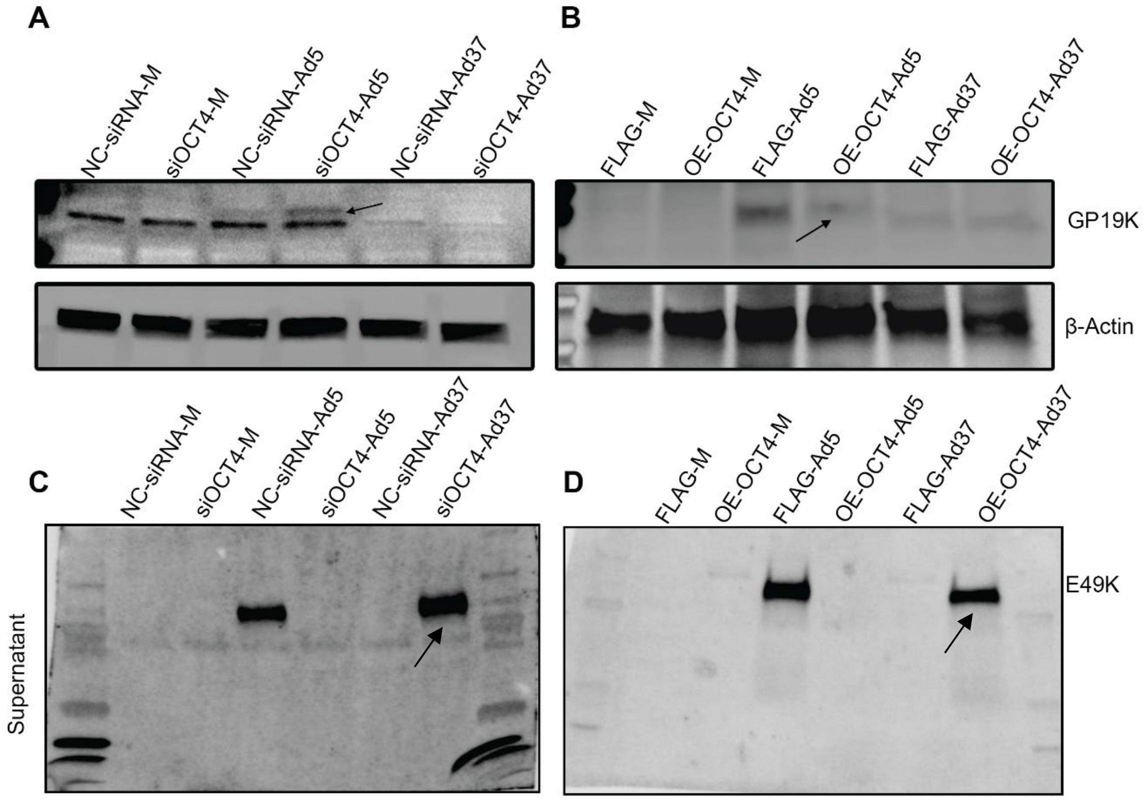

**Figure S5**. OCT4 regulates E3 protein expression. **(A)** OCT4-siRNA treated cells infected with HAdV-C5 show increased gp19k protein expression (arrow) compared to NC-siRNA treated cells. Note that the commercial gp19k antibody used cross reacted only with HAdV-C5 gp19k, not with HAdV-D37 gp19k. **(B)** In OCT4 overexpressing cells (OE-OCT4) infected with HAdV-C5 Gp19k expression was reduced (arrow)compared to HAdV-C5 infected Flag-only control cells (FLAG). **(C)** Supernatants collected from the same experiments as in A were probed with E49k antibody which recognizes HAdV-D37 protein, showing an increased secretion of E49K protein upon OCT4 knockdown as compared to NC-siRNA treated cells (arrows). **(D)** HAdV-D37- E49K showing reduced secretion in OE-OCT4 cells compared to FLAG control treated cells (arrows).

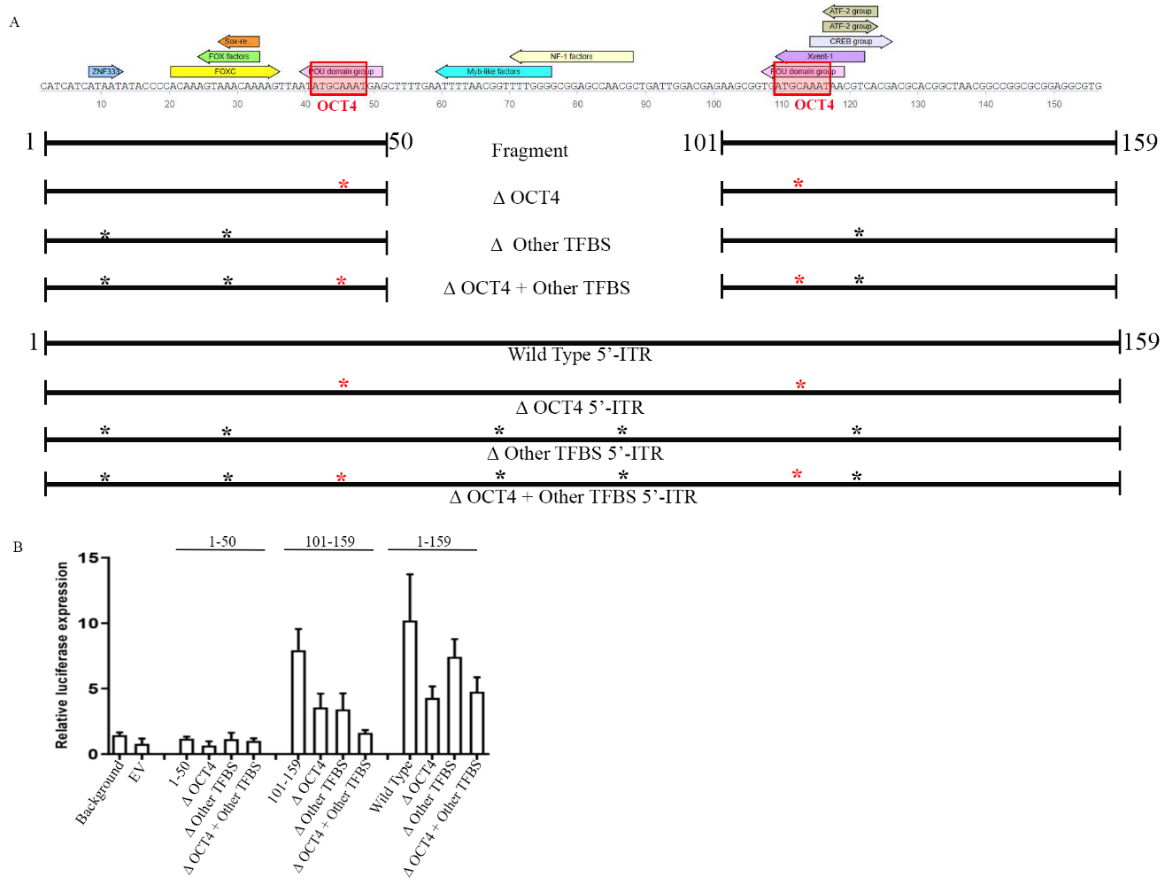

**Figure S6.** As compared to the OCT4 binding region within ntds 101-159, other putative transcription factor binding ITR sites (TFBS) show only relatively minor effects on downstream gene expression. **(A)** Schematic showing mutants generated for binding sites to OCT 4 (*) and other transcription factors (*). **(B)** Luciferase reporter assay showing reduction in promoter activity when both ATF2 and OCT4 binding sites were mutated in the 101-159 fragment. There was no significant reduction in the promoter activity of the full length ITR region encompassing ntds 1-159, when mutations of OCT4 and ATF2 binding sites were combined within the 5'-ITR.

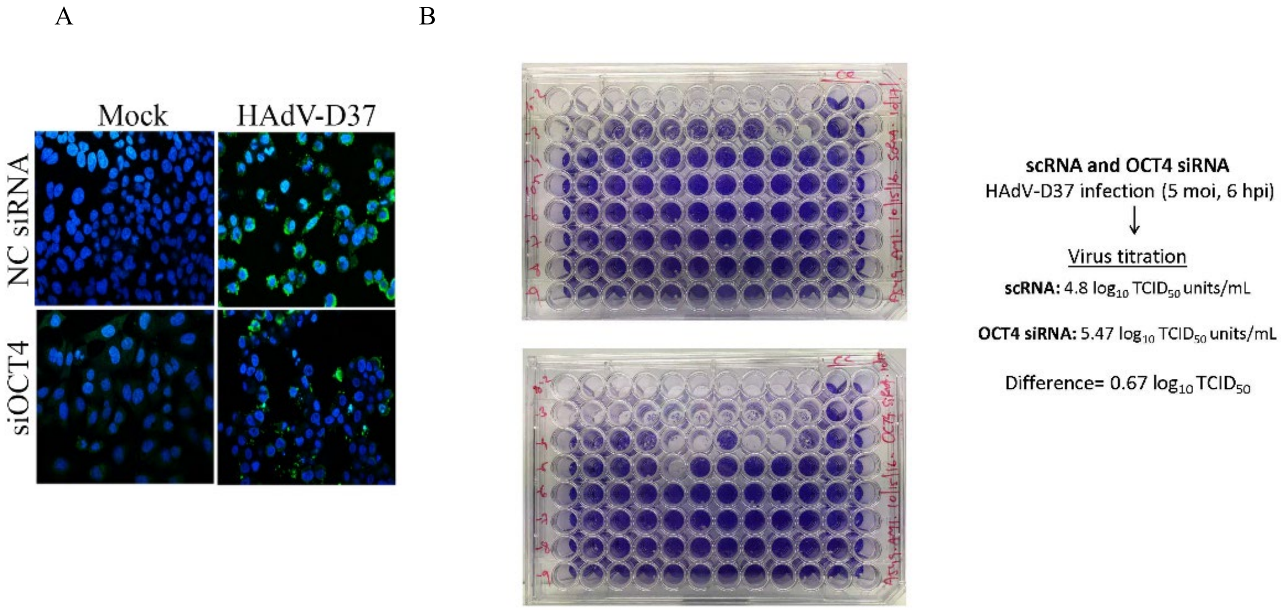

**Figure S7:** Apoptosis and HAdV-D37 replication. **(A)** TUNEL assay performed on control Nc-siRNA treated cells shows apoptosis (green) upon HAdV-D37 infection at 8hrs post infection. siOCT4 treated cells showed reduced apoptosis upon HAdV-D37 infection. **(B)** Supernatants collected after 6 hrs infection from OCT4 siRNA pretreated cells show increased viral titer by > ½ log ($TCID_{50}$) as compared to NC-SiRNA treated cells.

| Matrix name | Recognized factors |
|---|---|
| V$AHR_Q5 | AhR, AhR2, Ahr-xbb2 |
| V$AHRARNT_01 | AhR, AhR2, Ahr-xbb2, Arnt (774 AA form), arnt, arnt-L, arnt-S |
| V$AHRHIF_Q6 | AHH, AHH-isoform1, AHH-isoform2, AHH-isoform3, AhR, AhR repressor, AhR repressor:arnt, AhR2, AhR:arnt, Ahr-xbb2, Arnt (774 AA form), HIF-1, HIF-1alpha, HIF-1alpha-isoform1, HIF-1alpha-isoform2, HIF-1alpha-xbb2, arnt, arnt-L, arnt-S, arnt2 |
| V$AP1_Q2_01 | AP-1, FosB, Fra-1, Fra-2, Fra1, JunB, JunD, c-Jun, c-fos, deltaFosB, fra 1, jun B, jun D |
| V$AP1_Q4_01 | AP-1, FosB, Fra-1, Fra-2, Fra1, JunB, JunD, c-Jun, c-fos, deltaFosB, fra 1, jun B, jun D |
| V$AP2_Q6 | AP-2, AP-2alpha, AP-2alpha isoform 2, AP-2alpha isoform 3, AP-2alpha isoform 4, AP-2alphaA, AP-2alphaB, AP-2beta, AP-2gamma |
| V$AP2_Q6_01 | AP-2, AP-2alpha, AP-2alpha isoform 2, AP-2alpha isoform 3, AP-2alpha isoform 4, AP-2alphaA, AP-2alphaB, AP-2beta, AP-2gamma |
| V$AP2ALPHA_01 | AP-2, AP-2alpha, AP-2alpha isoform 2, AP-2alpha isoform 3, AP-2alpha isoform 4, AP-2alphaA, AP-2alphaB |
| V$AP4_01 | AP-4, TCFAP4, TFAP4 |
| V$AR_Q2 | AR, AR-isoform1, AR-isoform2 |
| V$AREB6_03 | ZEB, ZEB (1124 AA), ZEB (1154 AA), Zeb1, Zfhep-1, Zfhep-2 |
| V$ARNT_01 | Arnt (774 AA form), arnt, arnt-L, arnt-S |
| V$BACH2_01 | bach2 |
| V$BCL6_Q3 | Bcl-6 |
| V$BRCA_01 | brca1, brca1:USF2 |

| | |
|---|---|
| V$CACD_01 | CACD |
| V$CART1_01 | CART1 |
| V$CDP_02 | CDP, CDP-isoform1, CDP-isoform2, CDP-isoform4, CDP-isoform5, CDP-isoform6, CDP-isoform7, CDP2, NF-muNR |
| V$CDPCR1_01 | CDP, CDP-isoform1, CDP-isoform2, CDP-isoform4, CDP-isoform5, CDP-isoform6, CDP-isoform7, CDP2, NF-muNR |
| V$CDPCR3_01 | CDP, CDP-isoform1, CDP-isoform2, CDP-isoform4, CDP-isoform5, CDP-isoform6, CDP-isoform7, CDP2, NF-muNR |
| V$CDXA_02 | Cdx-1 |
| V$CEBP_Q2_01 | C/EBPalpha, C/EBPalpha(p20), C/EBPalpha(p30), C/EBPbeta, C/EBPbeta(LAP), C/EBPbeta(p20), C/EBPbeta(p34), C/EBPbeta(p35), C/EBPdelta, C/EBPgamma, LAP*-NF-M, NF-IL6-1, NF-IL6-2, NF-IL6-3, cebpe |
| V$CEBPA_01 | C/EBP, C/EBPalpha, C/EBPalpha(p20), C/EBPalpha(p30) |
| V$CEBPD_Q6 | C/EBPdelta |
| V$CEBPG_Q6 | C/EBPgamma |
| V$CETS1P54_01 | Ets-1 deltaVII, c-Ets-1, c-Ets-1 54, c-Ets-1A, c-Ets-1B |
| V$CETS1P54_02 | Ets-1 deltaVII, c-Ets-1, c-Ets-1 54, c-Ets-1A, c-Ets-1B |
| V$CETS1P54_03 | Ets-1 deltaVII, c-Ets-1, c-Ets-1 54, c-Ets-1A, c-Ets-1B |
| V$CIZ_01 | CIZ, CIZ-isoform1, CIZ-isoform2, CIZ5, CIZ6-1, CIZ6-2, CIZ6-3, CIZ8 |
| V$CKROX_Q2 | c-Krox |
| V$CMAF_01 | c-MAF, c-MAF-LONG, c-MAF-SHORT, c-MAF-isoform1, c-MAF-isoform2, mammary activating factor |
| V$COREBINDINGFACTOR_Q6 | core-binding factor |
| V$COUP_DR1_Q6 | COUP, COUP-TF II, COUP-TF1, COUP-TF2, COUP-TFI |
| V$CP2_02 | CP2, CP2 isoform 2, CP2-isoform1, CP2-isoform2, CP2-xbb2, Tcfcp2 |
| V$CREB_02 | CREB, CREBbeta, CREBomega, creb1, deltaCREB |

| | |
|---|---|
| V$CREB_Q3 | 120-kDa CRE-binding protein, 47-kDa CRE binding protein, ATF, ATF-1, ATF-2, ATF-2-isoform1, ATF-2-isoform2, ATF-2-isoform3, ATF-2-xbb3, ATF-2-xbb4, ATF-3, ATF-3 deltaZIP, ATF-3b, ATF-4, ATF-a, ATF4 (381AA), ATFa, ATFa-isoform1, ATFa-isoform2, ATFa-isoform3, Atf7, CREB, CREBbeta, CREBomega, CREM, CREM isoform-1, CREM isoform-2, CREM-Ia, CREM-Ib, CREMalpha, CREMbeta, CREMdeltaC-F, CREMdeltaC-G, CREMepsilon, CREMgamma, CREMtau, CREMtau1, CREMtau2, CREMtaualpha, ICER, ICER-I, ICER-II, ICER-IIgamma, ICER-Igamma, ICER-xbb1, S-CREM, S-CREMbeta, creb1, deltaCREB |
| V$CREBATF_Q6 | 120-kDa CRE-binding protein, 47-kDa CRE binding protein, ATF, ATF-1, ATF-2, ATF-2-isoform1, ATF-2-isoform2, ATF-2-isoform3, ATF-2-xbb3, ATF-2-xbb4, ATF-3, ATF-3 deltaZIP, ATF-3b, ATF-4, ATF-5, ATF-a, ATF-like, ATF/CREB, ATF4 (381AA), ATF6, ATFa, ATFa-isoform1, ATFa-isoform2, ATFa-isoform3, Atf5, Atf7, CREB, CREBbeta, CREBomega, CREM, CREM isoform-1, CREM isoform-2, CREM-Ia, CREM-Ib, CREMalpha, CREMbeta, CREMdeltaC-F, CREMdeltaC-G, CREMepsilon, CREMgamma, CREMtau, CREMtau1, CREMtau2, CREMtaualpha, ICER, ICER-I, ICER-II, ICER-IIgamma, ICER-Igamma, ICER-xbb1, S-CREM, S-CREMbeta, creb1, deltaCREB |
| V$CRX_Q4 | crx, rax |
| V$DBP_Q6 | DBP |
| V$DEAF1_02 | DEAF-1, DEAF1-3, DEAF1-4, Deaf1, Deaf1-2 |
| V$DEC_Q1 | DEC1, DEC2, Stra13 |
| V$DMRT3_01 | DMRT3 |

| | |
|---|---|
| V$DR1_Q3 | COUP, COUP-TF II, COUP-TF1, COUP-TF2, COUP-TFI, HNF gamma, HNF-4, HNF-4 alpha8, HNF-4alpha, HNF-4alpha1, HNF-4alpha2, HNF-4alpha3, HNF-4alpha4, HNF-4alpha7, HNF4A, HNF4gamma, NR1B1:RXR-alpha, PPARalpha:RXR-alpha, PPARgamma2:RXR-alpha, PPARgamma:RXR-alpha |
| V$DR3_Q4 | BXR-beta, CAR, CAR2:RXR-alpha, CAR:RXR-alpha, PXR, PXR-isoform1, PXR-isoform1:RXR-alpha, PXR-isoform1A, PXR-isoform1A:RXR-alpha, PXR-isoform1A:RXR-beta, PXR-isoform1B, PXR-isoform1C, PXR-isoform2, PXR-isoform2:RXR-alpha, PXR-isoform2A, PXR-isoform2B, PXR-isoform2C, PXR-isoform3, RXR-alpha, RXR-beta, SXR, SXR:RXR-alpha, VDR, VDR-M4 |
| V$DR4_Q2 | BXR-beta, CAR, CAR2:RXR-alpha, CAR:RXR-alpha, COUP, COUP-TF II, COUP-TF1, COUP-TF2, COUP-TFI, LXR-alpha, LXR-alpha:RXR-alpha, LXR-beta, LXR-beta:RXR-alpha, LXRalpha-isoform1, LXRalpha-isoform2, NR1B1, NR1B1-isoform1, NR1B1-isoform2, NR1B1:RXR-alpha, NR1B1:RXR-gamma, NR1B2, PXR, PXR-isoform1, PXR-isoform1:RXR-alpha, PXR-isoform1A, PXR-isoform1A:RXR-alpha, PXR-isoform1A:RXR-beta, PXR-isoform1B, PXR-isoform1C, PXR-isoform2, PXR-isoform2:RXR-alpha, PXR-isoform2A, PXR-isoform2B, PXR-isoform2C, PXR-isoform3, RAR-gamma, RARA, RXR-alpha, RXR-beta, RXR-beta1, RXR-beta2, SXR, SXR:RXR-alpha |
| V$E12_Q6 | E12, E2A, E47, ITF-1, Tcfe2a |
| V$E2_01 | E2 |
| V$E2_Q6_01 | E2 |

| | |
|---|---|
| V$E2A Q2 | E12, E2A, E47, ITF-1, MRF4, Myf-6, MyoD, MyoD (275 AA), MyoD (376 AA), Tcfe2a, myogenin |
| V$E2F 03 | E2F, E2F-1, E2F:DP |
| V$E2F Q6 01 | DP-1, E2F, E2F-1, E2F-1:DP-1, E2F-3, E2F-3B, E2F-3a, E2F-4, E2F-7, E2F:DP, E2F:DP:E4 |
| V$EBF Q6 | COE1, COE1-isoform1, COE1-long, COE1-short, COE1-xbb1, COE2, COE3, COE3-isoform1, COE3-isoform2 |
| V$EBOX Q6 01 | Arid5B, DEC1, DEC2, E12, E2A, E47, EHAND, EMF1, EMF2, EMF3, EMF4, HEB1-p67, HEB1-p94, HEN1, HTF4, HTF4alpha, HTF4beta, HTF4gamma, Hand2, INSAF, ITF, ITF-1, ITF-2, MASH-1, MASH-2, MEF1, MITF, MITF-2A, MITF-2B, MITF-2B-delta, MITF-A, MITF-A1, MITF-A2, MITF-B1, MITF-B2, MITF-C1, MITF-C2, MITF-H, MITF-H1, MITF-H2, MITF-H3, MITF-M, MITF-M1, MITF-M2, MRF2-isoform1, MRF2-isoform2, MRF4, Mad1, Mad3, Mad4, Max, Max-isoform1, Max-isoform2, Max-isoform3, Mrf2, Mrf2alpha, Mrf2beta, Mxi1, Myf-6, MyoD, MyoD (275 AA), MyoD (376 AA), N-Myc, N-Myc1, N-Myc2, SEF2-1A, SEF2-1B, SEF2-1D, Stra13, TFEA, TFEA-isoform1, TFEA-xbb1, TFEA-xbb2, TFEB-xbb1, Tal-1, Tal-2, Tal1-PP22, Tal1-PP39, Tal1-PP42, Tcfe2a, USF, USF1a, USF1b, USF2, USF2A-delta-H, USF2a, USF2b, USF2c, c-Myc, c-Myc I, c-Myc II, c-Myc-isoform1, c-Myc-isoform2, dHAND, myf-5, myogenin, tfeb, tfeb-isoform1, tfeb-isoform2, usf1, usf1:USF2, v-Myc |
| V$EFC Q6 | RFX1, RFX5:RFXAP:RFXANK |
| V$EGR1 01 | Egr-1 |

| | |
|---|---|
| V$ER_Q6 | ER-alpha, ER-alpha-L, ER-alpha-S |
| V$ETF_Q6 | ETF |
| V$ETS_Q6 | ELFR, ETV7, Elf-1, Elf4, Elk-1, Elk1-isoform1, Elk1-isoform2, Erg, Erg isoform 1, Erg isoform 2, Erg isoform 3, Erg isoform 5, Erg-1, Erg-2, Ets-1 deltaVII, Fli-1, GABP-alpha, GABP-alpha:GABP-beta, GABP-beta, GABP-beta1, GABP-beta2, GABP-gamma1, GABP-gamma2, NRF-2beta1, NRF-2gamma1, Net, Net-b, PEA3, PU.1-xbb1, SAP-1, SAP-1a, SAP-1b, SAP1A, SPI1, Spi-B, Spi-B-IA, Spi-B-IIA, Spi-B-isoform1, Spi-B-isoform2, TCF, TEL1, Tel-2a, Tel-2b, Tel-2c, Tel-2d, Tel-2e, Tel-2f, c-Ets-1, c-Ets-1 54, c-Ets-1 68, c-Ets-1A, c-Ets-1B, c-Ets-2, c-Ets-2 58-64, c-Ets-2A, c-Ets-2B, nerf, nerf-isoform1, nerf-isoform2, nerf-isoform3, nerf-isoform4, nerf-isoform5, p38erg, p49erg, p55, p55erg |
| V$FAC1_01 | FAC1, FAC1-xbb1, Falz |
| V$FOXJ2_02 | FOXJ2 (long isoform), FOXJ2 (short isoform), Foxj2 |
| V$FXR_Q3 | FXR, FXR-alpha1, FXR-alpha2, FXR-beta1, FXR-beta2, FXR-isoform1, FXR-isoform2, FXR-isoform3, FXR-isoform3:RXR-alpha, FXR-isoform4, FXR:RXR-alpha |
| V$GATA_C | GATA-1, GATA-1 isoform 1, GATA-1 isoform 2, GATA-1A, GATA-1B, GATA-2, GATA-3, GATA-3 isoform-1, GATA-3 isoform-2 |
| V$GATA_Q6 | GATA-1, GATA-1 isoform 1, GATA-1 isoform 2, GATA-1A, GATA-1B, GATA-2, GATA-3, GATA-3 isoform-1, GATA-3 isoform-2, GATA-4, GATA-5, GATA-5A, GATA-5B, GATA-6, GATA-6A, GATA-6B, GATA-6long, GATA-6short |

| | |
|---|---|
| V$GCNF_01 | GCNF, GCNF-1, GCNF-2 |
| V$GEN_INI3_B | |
| V$GFI1_Q6 | Gfi1, gfi1b |
| V$GLI_Q2 | GLI, GLI2alpha, GLIS1, Gli2, Gli2beta, Gli2delta, Gli2gamma, GliH1, XGli3, gli3 |
| V$GRE_C | GR, GR-2, GR-Abeta, GR-Alpha-2, GR-Beta-2, GR-alpha, GR-beta |
| V$HIC1_03 | HIC-1, HIC-1-isoform1, HIC-1-isoform2 |
| V$HIF1_Q3 | HIF-1, HIF-1alpha, HIF-1alpha-isoform1, HIF-1alpha-isoform2, HIF-1alpha-xbb2 |
| V$HLF_01 | HLF17, HLF24, HLF36, HLF43, Hlf |
| V$HMGIY_Q6 | HMGI-C, HMGIY, HMGIY-isoform1, HMGIY-isoform2, HMGIY-isoform3 |
| V$HNF1_Q6 | HNF-1, HNF-1A, HNF-1alpha, HNF-1alpha-A, HNF-1alpha-B, HNF-1alpha-C, HNF-1beta, HNF-1beta-1, HNF-1beta-2, HNF-1beta-3, HNF-1beta-4, HNF-1beta-A, HNF-1beta-B, HNF-1beta-C, tcf2 |
| V$HNF3_Q6_01 | FoxA1, HNF-3, HNF-3beta, HNF-3gamma, HNF3(-like), HNF3A |
| V$HNF3B_01 | HNF-3beta |
| V$HNF4_Q6_01 | HNF-4, HNF-4 alpha8, HNF-4alpha, HNF-4alpha1, HNF-4alpha2, HNF-4alpha3, HNF-4alpha4, HNF-4alpha7, HNF4A |
| V$HNF4ALPHA_Q6 | HNF-4, HNF-4 alpha8, HNF-4alpha, HNF-4alpha1, HNF-4alpha2, HNF-4alpha3, HNF-4alpha4, HNF-4alpha7, HNF4A |
| V$HNF6_Q6 | HNF-6alpha, HNF-6beta, HNF6, HNF6-alpha, HNF6-beta, OC-2 |
| V$HOXA7_01 | HOXA7 |
| V$HSF1_Q6 | HSF1, HSF1-L, HSF1-S, HSF1long, HSF1short |
| V$HSF2_01 | HSF2, HSF2A, HSF2B |
| V$IK_Q5 | Ikaros |
| V$IPF1_Q4_01 | ipf1, ipf1:Pbx |

| | |
|---|---|
| V$IRF2_01 | IRF-2 |
| V$IRF_Q6 | IRF-1, IRF-10, IRF-2, IRF-3, IRF-4, IRF-5, IRF-6, IRF-7, IRF-7A, IRF-7B, IRF-7H, IRF-8, IRF4-1, IRF6 |
| V$ISRE_01 | ISGF-3 |
| V$KAISO_01 | Kaiso |
| V$KID3_01 | Kid3, Zfp354c |
| V$KROX_Q6 | Egr-1, Egr-2, Egr-4, Egr4, Krox20, Krox20-isoform1, Krox20-isoform2, egr3 |
| V$LEF1_Q2_01 | LEF-1, LEF-1B, LEF-1S, LEF1-isoform1, LEF1-isoform2, LEF1-isoform3, LEF1-isoform4 |
| V$LMO2COM_01 | RBTN2, RBTN2-isoform1 |
| V$LRF_Q2 | LRF, LRF-isoform1, LRF-isoform2 |
| V$LRH1_Q5 | LRH-1, LRH-1-xbb1, LRH1-isoform1, LRH1-isoform2, LRH1-isoform3 |
| V$LUN1_01 | LUN, LUN-isoform1, LUN-isoform2 |
| V$LXR_Q3 | LXR-alpha, LXR-alpha:RXR-alpha, LXR-beta, LXR-beta:RXR-alpha, LXRalpha-isoform1, LXRalpha-isoform2 |
| V$MAF_Q6_01 | Bach1, Bach1:MafK, Bach1t, MafB, MafF, MafG, MafG:MafG, MafK, NF-E2, NF-E2 p45, NF-E2p45, Nfe2l1, Nfe2l1-isoform1, Nfe2l1-isoform2, Nfe2l1-isoform2:MafG, Nfe2l1-isoform2:MafK, Nfe2l1-long, Nfe2l1-short, Nfe2l1-xbb1, Nrf2, Nrf2-isoform1, Nrf2-isoform2, Nrf2:MafG, Nrf2:MafK, Nrf3, Nrf3:MafK, bach2, c-MAF, c-MAF-LONG, c-MAF-SHORT, c-MAF-isoform1, c-MAF-isoform2, mammary activating factor, v-Maf |
| V$MAZ_Q6 | MAZ, MAZi, SAF-1, SAF-2, SAF1 |

| | |
|---|---|
| V$MEF2_Q6_01 | MEF-2A, MEF-2C, MEF-2C/delta32, MEF-2C/delta8, MEF-2C/delta8,32, MEF-2DA'0, MEF-2DA0, MEF-2DAB, MEF2A-xbb5, MEF2C, MEF2C isoform 1, MEF2C isoform 3, MEF2C isoform 4, MEF2C isoform 5, RSRFC4, RSRFC9, aMEF-2, mef2a, mef2a-isoform1, mef2a-isoform2, mef2a-isoform3, mef2a-isoform6, mef2d, mef2d-isoform1, mef2d-isoform2, mef2d-isoform3, mef2d-isoform4, mef2d-isoform5 |
| V$MEIS1_01 | Meis-1-1, Meis-1-2, Meis-1-3, meis1, meis1-a, meis1-b, meis1-c, meis1a, meis1b |
| V$MEIS1BHOXA9_02 | hoxa-9, hoxa-9T, hoxa9, hoxa9A, hoxa9B, meis1, meis1-a, meis1-b, meis1-c, meis1a, meis1b |
| V$MINI19_B | |
| V$MRF2_01 | Arid5B, MRF2-isoform1, MRF2-isoform2, Mrf2, Mrf2alpha, Mrf2beta |
| V$MTF1_Q4 | MTF-1 |
| V$MYB_Q3 | c-Myb, c-Myb-isoform1, c-Myb-isoform2, c-Myb-isoform3, c-Myb-isoform4, c-Myb-isoform5, c-Myb-isoform6 |
| V$MYCMAX_03 | Max, Max-isoform1, Max-isoform2, Max-isoform3, c-Myc, c-Myc I, c-Myc-isoform1, c-Myc-isoform2 |
| V$MYOGNF1_01 | CTF-1, CTF-2, CTF-3, CTF-5, CTF-7, NF-1, NF-1/L, NF-1/Red1, NF-1A, NF-1A1, NF-1A2, NF-1A3, NF-1A4, NF-1B, NF-1B-xbb5, NF-1B1, NF-1B2, NF-1B3, NF-1B4, NF-1C, NF-1C1, NF-1C2, NF-1C4, NF-IX, NF1C-isoform1, NFI, NFIA1-xbb1, NFIA1-xbb2, NFIA2, NFIA2.1, NFIB1, NFIB2, NFIB3, NFIB6, NFIC10, NFIC11, NFIC1A, NFIC1B, NFIC2, NFIC5, NFIC8, NFIC9 |

| | |
|---|---|
| V$NF1_Q6_01 | CTF-1, CTF-2, CTF-3, CTF-5, CTF-7, NF-1, NF-1/L, NF-1A, NF-1A1, NF-1A2, NF-1A3, NF-1A4, NF-1C, NF-1C1, NF-1C2, NF-1C4, NF1C-isoform1, NFI, NFIA1-xbb1, NFIA1-xbb2, NFIA2, NFIA2.1, NFIB6, NFIC10, NFIC11, NFIC1A, NFIC1B, NFIC2, NFIC5, NFIC8, NFIC9 |
| V$NFAT_Q4_01 | NF-AT1, NF-AT1B, NF-AT1C, NF-AT2, NF-AT3, NF-AT4, NFAT1, NFAT1-isoformB, NFAT1-isoformC, NFAT1-isoformD |
| V$NFKB_Q6_01 | NF-TNF, NF-kappaB, NF-kappaB(-like), NF-kappaB1, NF-kappaB1-isoform1, NF-kappaB1-isoform2, NF-kappaB1-isoform3, NF-kappaB1-isoform4, NF-kappaB1-isoform5, NF-kappaB1-isoform6, NF-kappaB1-isoform7, NF-kappaB1-p50, NF-kappaB2, NF-kappaB2 (p49), NF-kappaB2-p100, NF-kappaB2-p52, NFKappaB1, NFkappaB2, Rel-A-p65, RelA-p65, RelA-p65 delta 2, RelA-p65delta, p100, p105, p50, p65 delta |
| V$NFY_01 | CBF(2), NF-Y |
| V$NFY_Q6_01 | CBF(2), CBF-B, CBF-C, CP1, NF-Y, NF-YA, NF-YA isoform-1, NF-YA isoform-2, NF-YB, NF-YC, NF-YC-1, NF-YC-3, NF-Yprime, NFY-A |
| V$NKX22_01 | NKX2B |
| V$NKX25_Q5 | CSX |
| V$NKX3A_01 | NKX3A, NKX3A-V1, NKX3A-V2, NKX3A-V3, NKX3A-V4 |
| V$NRSF_Q4 | REST, REST isoform 1, REST4, RESTisoform3, RESTisoform4, Rest isoform1, Rest isoform2 |
| V$OCT1_02 | POU2F1, POU2F1-isoform1, POU2F1-isoform2, POU2F1a, POU2F1b, POU2F1c |
| V$OCT1_03 | POU2F1, POU2F1-isoform1, POU2F1-isoform2, POU2F1a, POU2F1b, POU2F1c |

| | |
|---|---|
| V$OCT1_07 | POU2F1, POU2F1-isoform1, POU2F1-isoform2, POU2F1a, POU2F1b, POU2F1c |
| V$OCT1_Q5_01 | POU2F1, POU2F1-isoform1, POU2F1-isoform2, POU2F1a, POU2F1b, POU2F1c |
| V$OCT4_01 | Oct3, Oct3-isoform1, Oct3-isoform2, Oct3-xbb1, POU5F1 (Oct-5) |
| V$OCT4_02 | Oct3, Oct3-isoform1, Oct3-isoform2, Oct3-xbb1, POU5F1 (Oct-5) |
| V$OCT_Q6 | BRN1, Brn-3a, OCA-B, OTF3P1, Oct-10, Oct-2, Oct-2.1, Oct-2.2, Oct-2.3, Oct-2.4, Oct-2.5, Oct-2.6, Oct-2B, Oct-9, Oct-R, Oct3, Oct3-isoform1, Oct3-isoform2, Oct3-xbb1, Octa-factor, Octamer binding factor, POU2F1, POU2F1-isoform1, POU2F1-isoform2, POU2F1a, POU2F1b, POU2F1c, POU2F2, POU2F2 (Oct-2.1), POU2F2 (Oct-2.7), POU2F2 (Oct-2.8), POU2F2B, POU2F3, POU3F1, POU3F2, POU3F2 (N-Oct-5a), POU3F2 (N-Oct-5b), POU4F1, POU4F1(l), POU5F1 (Oct-5), oct-B2, oct-B3 |
| V$OG2_01 | OG-2 |
| V$OSF2_Q6 | AML3, AML3-G1, AML3-G2, AML3-U1, AML3-Y1, AML3-Y2, AML3-isoform1, AML3-isoform2, AML3-isoform3, PEBP2alphaA1, PEBP2alphaA2 |
| V$P300_01 | p300 |
| V$P53_02 | Delta113p53, Delta133p53, Delta133p53beta, Delta133p53gamma, Delta40p53, Delta40p53beta, Delta40p53gamma, p53, p53-isoform1, p53as, p53beta, p53gamma |
| V$PAX2_01 | Pax-2a, Pax-2b, pax2, pax2-isoform1, pax2-isoform2, pax2-isoform3 |
| V$PAX3_01 | PAX3, Pax-3 |
| V$PAX4_03 | Pax-4a, Pax4 |
| V$PAX4_04 | Pax-4a, Pax4 |
| V$PAX5_01 | Pax-5 |

| | |
|---|---|
| V$PAX5_02 | Pax-5 |
| V$PAX6_Q2 | Pax-6 (Pax-QNR), pax6, pax6-isoform1, pax6-isoform2, pax6-isoform5a |
| V$PAX8_01 | Pax-8, Pax-8a, Pax-8c, Pax-8d, Pax-8e, Pax-8f |
| V$PBX1_02 | Pbx1, Pbx1a, Pbx1b |
| V$PBX_Q3 | PBX3c, Pbx, Pbx1, Pbx1:HOXB1, Pbx1:Prep1, Pbx1a, Pbx1a:HOXB7, Pbx1a:HOXB8, Pbx1a:HOXC6, Pbx1a:PREP1, Pbx1a:ipf1, Pbx1b, Pbx1b:Prep1, Pbx2, Pbx2:HOXC6, Pbx2:Prep1, Pbx3, Pbx3a, Pbx3a:HOXC6, Pbx3a:PREP1, Pbx3b |
| V$PLZF_02 | PLZFA, PLZFB, Plzf |
| V$POU1F1_Q6 | POU1F1, Pit-1, Pit-1-xbb3, Pit-1A, Pit-1B, Pit-1beta |
| V$POU3F2_01 | POU3F2, POU3F2 (N-Oct-5a), POU3F2 (N-Oct-5b) |
| V$POU3F2_02 | POU3F2, POU3F2 (N-Oct-5a), POU3F2 (N-Oct-5b) |
| V$POU6F1_01 | POU6F1, POU6F1 (c1), POU6F1 (c2), POU6F1 (c7) |
| V$PPAR_DR1_Q2 | ORF4-gamma, PPARalpha, PPARalpha:RXR-alpha, PPARbeta, PPARdelta, PPARdelta-isoform1, PPARdelta-isoform2, PPARgamma, PPARgamma1, PPARgamma2, PPARgamma2:RXR-alpha, PPARgamma:RXR-alpha |
| V$PPARA_01 | PPARalpha |
| V$PPARA_02 | PPARalpha, RXR-alpha |
| V$PPARG_01 | ORF4-gamma, PPARgamma, PPARgamma1, PPARgamma2 |
| V$PPARG_02 | ORF4-gamma, PPARgamma, PPARgamma1, PPARgamma2 |
| V$R_01 | R |

| | |
|---|---|
| V$RBPJK_Q4 | RBPJK, RBPJK-isoform1, RBPJK-isoform2, RBPJK-isoform3, RBPJK-isoform4, RBPJK-isoform5, RBPJK-isoform6 |
| V$RFX_Q6 | RFX-B-delta5, RFX1, RFX1:RFX3, RFX1:rfx2, RFX3, RFX5, RFX5:RFXAP:RFXANK, RFXANK, RFXANK short isoform, RFXAP, rfx2, rfx4, rfx4-isoform1, rfx4-isoform2, rfx4-isoform3, rfx4-isoform4 |
| V$RORA_Q4 | RORalpha, RORalpha-4, RORalpha1, RORalpha2, RORalpha3 |
| V$RP58_01 | RP58 |
| V$RSRFC4_Q2 | MEF-2A, MEF2A-xbb5, RSRFC4, RSRFC9, aMEF-2, mef2a, mef2a-isoform1, mef2a-isoform2, mef2a-isoform3 |
| V$RUSH1A_02 | HLTF (Met123), RUSH-1, RUSH-1alpha, RUSH-1beta, ZBU1 |
| V$SF1_Q6 | SF-1, SF1, SF1- isoform1, SF1- isoform2, SF1-Xbb1 |
| V$SMAD3_Q6 | Smad3 |
| V$SOX9_B1 | Sox9 |
| V$SP1_Q6 | Sp1, Sp1-isoform1, Sp1-isoform2 |
| V$SP3_Q3 | Sp3-isoform1, Sp3-isoform2, Sp3-isoform3, Sp3-isoform4, Sp3-isoform5, Sp3-isoform6, Sp3-xbb1, sp3 |
| V$SPZ1_01 | SPZ1 isoform1, Spz1 |
| V$SREBP1_01 | SREBP-1, SREBP-1a, SREBP-1b, SREBP-1c |
| V$SREBP_Q3 | SREBP-1, SREBP-1a, SREBP-1b, SREBP-1c, SREBP-2 |
| V$SREBP_Q6 | SREBP-1, SREBP-1a, SREBP-1b, SREBP-1c, SREBP-2 |

| | |
|---|---|
| V$SRF_C | SRF, SRF-I, SRF-L, SRF-M, SRF-S |
| V$SRF_Q6 | SRF, SRF-I, SRF-L, SRF-M, SRF-S |
| V$STAF_02 | Staf, Zfp143 |
| V$STAT1_01 | STAT1, STAT1alpha, STAT1beta |
| V$STAT_Q6 | STAT1, STAT1alpha, STAT1beta, STAT2, STAT3, STAT3-isoform1, STAT3-isoform2, STAT3-isoform3, STAT4, STAT5A, STAT5B, STAT6, STAT6-isoform1, STAT6-isoform2, Stat5A1, Stat5A2 |
| V$SZF11_01 | SZF1, SZF1-isoform2, SZF1-isoform3 |
| V$TAL1BETAE47_01 | E12, E2A, E47, ITF-1, Tal-1, Tal1-PP22, Tal1-PP39, Tal1-PP42, Tcfe2a |
| V$TAXCREB_01 | CREB, CREBbeta, CREBomega, Tax, creb1, deltaCREB |
| V$TAXCREB_02 | CREB, CREBbeta, CREBomega, Tax, creb1, deltaCREB |
| V$TBP_Q6 | TBP, TFIID |
| V$TBX5_01 | TBX5-L, TBX5-S, Tbx5 |
| V$TCF11_01 | Nfe2l1, Nfe2l1-isoform1, Nfe2l1-isoform2, Nfe2l1-long, Nfe2l1-short, Nfe2l1-xbb1 |
| V$TCF4_Q5 | TCF 4, TCF-4, TCF-4(K), TCF-4-isoform1, TCF-4-isoform10, TCF-4-isoform2, TCF-4-isoform3, TCF-4-isoform4, TCF-4-isoform5, TCF-4-isoform6, TCF-4-isoform7, TCF-4-isoform8, TCF-4-isoform9, TCF-4B, TCF-4E, TCF4 |
| V$TEL2_Q6 | ETV7, Tel-2a, Tel-2b, Tel-2c, Tel-2d, Tel-2e, Tel-2f |
| V$TFE_Q6 | MITF, MITF-A, MITF-A1, MITF-A2, MITF-B1, MITF-B2, MITF-C1, MITF-C2, MITF-H, MITF-H1, MITF-H2, MITF-H3, MITF-M, MITF-M1, MITF-M2, TFEA, TFEA-isoform1, TFEA-xbb1, TFEA-xbb2, TFEB-xbb1, tcfec, tfeb, tfeb-isoform1, tfeb-isoform2 |
| V$TST1_01 | POU3F1 |

| | |
|---|---|
| V$USF2_Q6 | USF2, USF2A-delta-H, USF2a, USF2b, USF2c, usf1:USF2 |
| V$VDR_Q3 | VDR, VDR-M4 |
| V$VJUN_01 | v-Jun |
| V$VMYB_02 | v-Myb |
| V$WT1_Q6 | WT1, WT1 -KTS, WT1 I, WT1 I -KTS, WT1 I-del2, WT1-del2, WT1-isoform1, WT1-isoform2, WT1-isoform3, WT1-isoform4 |
| V$XFD2_01 | FOXI1a |
| V$XVENT1_01 | Xvent-1 |
| V$YY1_01 | NF-E, YY1 |
| V$YY1_Q6 | NF-E, YY1 |
| V$ZF5_B | ZF5, ZFP161 |
| V$ZIC3_01 | Zic3 |
| V$ZID_01 | ZID |

| Species | Number of sites | core similarity score (CSS) | Matrix similarity score (MSS) |
|---|---|---|---|
| guinea pig, human, killifish, mouse, pig, rabbit, rat | 25 | 0.7 | 0.946 |
| Mammalia, guinea pig, human, killifish, mouse, pig, rabbit, rat | 24 | 0.721 | 0.89 |
| Mammalia, guinea pig, human, killifish, mouse, pig, rabbit, rat | 47 | 0.752 | 0.979 |
| Japanese pufferfish, Japanese quail, Mammalia, cattle, chick, dog, domestic pig, hamster, human, monkey, mouse, rabbit, rat, sheep | 46 | 0.822 | 0.951 |
| Japanese pufferfish, Japanese quail, Mammalia, cattle, chick, dog, domestic pig, hamster, human, monkey, mouse, rabbit, rat, sheep | 119 | 0.818 | 0.957 |
| Chinese hamster (Gray hamster), Mammalia, clawed frog, human, mouse, rabbit, rat | 13 | 0.743 | 0.844 |
| Chinese hamster (Gray hamster), Mammalia, clawed frog, human, mouse, rabbit, rat | 24 | 0.684 | 0.9 |
| Chinese hamster (Gray hamster), Mammalia, human, mouse, rabbit, rat | 185 | 0.83 | 0.981 |
| cattle, human, mouse, rat | 5 | 0.679 | 0.834 |
| Mammalia, Vertebrata, baboon, bull frog, chimpanzee, dog, human, lemur, macaque, mouse, pig, rabbit, rat, spotted hyena | 7 | 0.719 | 0.777 |
| Mammalia, chick, golden (Syrian) hamster, human, mouse, rat | 12 | 0.62 | 0.906 |
| Mammalia, human, mouse, rat | 20 | 0.712 | 0.835 |
| human, mouse, rat | 26 | 0.734 | 0.963 |
| Mammalia, human, mouse, rat | 7 | 0.718 | 0.9 |
| Mammalia, human, rat | 43 | 0.74 | 0.955 |

| | | | |
|---|---|---|---|
| mouse | 17 | 0.601 | 0.841 |
| clawed frog, human, mouse, rat | 25 | 0.773 | 0.871 |
| Mammalia, human, mouse, rat | 86 | 0.613 | 0.756 |
| Mammalia, human, mouse, rat | 32 | 0.72 | 0.777 |
| Mammalia, human, mouse, rat | 24 | 0.632 | 0.737 |
| chick | 18 | 0.843 | 0.925 |
| Mammalia, cattle, chick, human, mouse, pig, rat | 89 | 0.758 | 0.921 |
| Mammalia, chick, clawed frog, human, mouse, rat | 43 | 0.887 | 0.92 |
| Mammalia, human, mouse, rat | 16 | 0.796 | 0.911 |
| Mammalia, human, mouse, rat | 7 | 0.644 | 0.777 |
| Mammalia, chick, clawed frog, human, mouse, rat | 15 | 0.75 | 0.928 |
| Mammalia, chick, clawed frog, human, mouse, rat | 40 | 0.901 | 0.948 |
| Mammalia, chick, clawed frog, human, mouse, rat | 33 | 0.833 | 0.892 |
| human, mouse, rat | 39 | 0.9 | 0.996 |
| Mammalia, human, mouse, rabbit | 10 | 0.765 | 0.938 |
| Mammalia, chick, human, mouse, rat | 37 | 0.65 | 0.836 |
| cat, cattle, mouse | 9 | 0.702 | 0.864 |
| Mammalia, cattle, chick, clawed frog, human, monkey, mouse, pig, rat | 9 | 0.567 | 0.746 |
| Mammalia, human, mouse, rat | 19 | 0.681 | 0.76 |
| Mammalia, cattle, domestic pig, hamster, human, mink, monkey, mouse, rabbit, rat, sheep | 16 | 0.717 | 0.845 |

| | | | |
|---|---|---|---|
| Mammalia, cattle, domestic pig, hamster, human, mink, monkey, mouse, rabbit, rat, sheep | 53 | 0.735 | 0.953 |
| Mammalia, cattle, clawed frog, domestic pig, gibbon ape, hamster, human, mink, monkey, mouse, rabbit, rat, sheep | 60 | 0.73 | 0.895 |
| cattle, clawed frog, human, mouse, rat | 26 | 0.902 | 0.925 |
| Mammalia, human, mouse, rat | 15 | 0.778 | 0.959 |
| Mammalia, chimpanzee, human, mouse, rat | 21 | 0.657 | 0.824 |
| Mammalia, human, mouse, rat | 5 | 0.874 | 0.934 |
| human, mouse, rat, zebra fish | 40 | 0.734 | 0.881 |

| | | | |
|---|---|---|---|
| Mammalia, cattle, chick, chipmunk, clawed frog, human, monkey, mouse, pig, rat | 31 | 0.675 | 0.812 |
| Mammalia, chick, clawed frog, domestic pig, human, monkey, mouse, quail, rat | 14 | 0.7 | 0.734 |
| Mammalia, cattle, chick, clawed frog, human, monkey, mouse, pig, quail, rat | 19 | 0.545 | 0.7 |
| Chinese hamster (Gray hamster), Mammalia, clawed frog, golden (Syrian) hamster, human, mouse, rat | 11 | 0.74 | 0.915 |
| BPV-1 | 17 | 0.795 | 0.952 |
| BPV-1 | 18 | 0.756 | 0.942 |

| | | | |
|---|---|---|---|
| Chinese hamster (Gray hamster), Mammalia, chick, clawed frog, common carp, golden (Syrian) hamster, human, monkey, mouse, quail, rainbow trout, rat, zebra fish | 13 | 0.846 | 0.916 |
| Mammalia, chick, human, mouse, rat | 29 | 0.644 | 0.811 |
| Mammalia, chick, human, mouse, rat | 39 | 0.726 | 0.817 |
| Mammalia, human, mouse, rat | 5 | 0.616 | 0.871 |
| Avian myelocytomatosis virus CMII, Avian myelocytomatosis virus MC29, Chinese hamster (Gray hamster), Feline leukemia provirus FTT, HBI, Kenyan clawed frog, MH2E21, Mammalia, OK10, canary, cattle, chick, chimpanzee, clawed frog, common carp, common marmoset, dog, domestic pig, duck, gibbon ape, golden (Syrian) hamster, goldfish, hamster, human, monkey, mouse, pig, quail, rabbit, rainbow trout, rat, sheep, wild cat, woodchuck, zebra fish | 119 | 0.683 | 0.898 |
| Mammalia, human, mouse, rat | 6 | 0.635 | 0.75 |
| Chinese hamster (Gray hamster), Mammalia, cattle, human, monkey, mouse, pig, rabbit, rat | 0 | 0.75 | 0.778 |

| | | | |
|---|---|---|---|
| Israeli tilapia, Mammalia, Nile tilapia, atlantic croaker, atlantic salmon, cattle, chick, clawed frog, desert grassland whiptail lizard, gilthead sea bream, hamster, horse, human, medaka, mouse, pig, rabbit, rat, red sea bream, sheep, zebra finch, zebra fish | 20 | 0.719 | 0.901 |
| human, rat | 13 | 0.854 | 0.989 |
| Chinese hamster (Gray hamster), Mammalia, chick, clawed frog, domestic pig, human, mouse, rat, zebra fish | 81 | 0.827 | 0.942 |
| Mammalia, human, mouse | 24 | 0.746 | 0.866 |
| human, mouse, rat | 6 | 0.578 | 0.772 |
| Mammalia, cattle, golden (Syrian) hamster, human, mouse, rat | 6 | 0.659 | 0.757 |
| Mammalia, cat, cattle, chick, clawed frog, human, mouse, rat | 0 | 0.707 | 0.905 |
| Mammalia, cat, cattle, chick, clawed frog, human, mouse, pig, quail, rabbit, rat | 105 | 0.808 | 0.974 |

| | | | |
|---|---|---|---|
| Mammalia, human, mouse | 37 | 0.666 | 0.85 |
| | 20 | 0.82 | 0.962 |
| Mammalia, human, mouse, rat | 25 | 0.835 | 0.968 |
| Mammalia, clawed frog, human, mouse | 15 | 0.735 | 0.88 |
| Bolivian squirrel monkey, Japanese flounder, Ma's night monkey, Mammalia, cat, chick, clawed frog, common squirrel monkey, common tree shrew, cotton-top tamarin, guinea pig, human, mouse, pig, rabbit, rainbow trout, rat, sheep | 0 | 0.619 | 0.702 |
| human, mouse | 32 | 0.937 | 0.936 |
| Mammalia, cattle, human, mouse, rat | 23 | 0.711 | 0.882 |
| human, mouse, rat | 18 | 0.743 | 0.902 |
| Mammalia, human, mouse, rat | 13 | 0.739 | 0.913 |
| Mammalia, chick, domestic pig, golden (Syrian) hamster, hamster, human, mouse, pig, rat | 34 | 0.723 | 0.864 |
| Mammalia, atlantic salmon, chick, clawed frog, golden (Syrian) hamster, hamster, human, mouse, rat, zebra fish | 63 | 0.874 | 0.939 |
| Mammalia, chick, clawed frog, human, mouse, rat | 24 | 0.835 | 0.906 |
| Mammalia, cattle, chipmunk, clawed frog, human, mouse, rat | 58 | 0.679 | 0.838 |
| Mammalia, cattle, chipmunk, clawed frog, human, mouse, rat | 11 | 0.688 | 0.8 |
| Mammalia, golden (Syrian) hamster, hamster, human, mouse, rat | 13 | 0.717 | 0.864 |
| clawed frog, human, mouse, quail | 18 | 0.9 | 0.998 |
| Chinese hamster (Gray hamster), Mammalia, chick, human, monkey, mouse, rat | 11 | 0.8 | 0.857 |
| chick, human, mouse, rat | 27 | 0.702 | 0.907 |
| Mammalia, human, mouse, rat | 21 | 0.733 | 0.831 |
| Mammalia, clawed frog, golden (Syrian) hamster, hamster, human, mouse, rat | 16 | 0.738 | 0.891 |

| | | | |
|---|---|---|---|
| Mammalia, human, mouse, rat, sheep | 15 | 0.575 | 0.864 |
| Mammalia, cattle, chick, clawed frog, human, mouse, rat, sheep | 26 | 0.886 | 0.965 |
| human | 13 | 0.7 | 0.848 |
| human, mouse | 32 | 0.7 | 0.91 |
| human, mouse | 16 | 0.871 | 0.997 |
| Chinese hamster (Gray hamster), Mammalia, cattle, human, monkey, mouse, pig, rabbit, rat | 23 | 0.776 | 0.887 |
| Mammalia, chick, clawed frog, human, mouse, rat, zebra fish | 23 | 0.799 | 0.957 |
| Mammalia, human, mouse, rat | 31 | 0.825 | 0.971 |
| Mammalia, human, mouse, rat | 11 | 0.864 | 0.968 |
| Mammalia, chick, human, mouse, rat | 14 | 0.734 | 0.93 |
| human | 24 | 0.9 | 0.684 |
| Mammalia, human, mouse, rat | 8 | 0.59 | 0.751 |
| AS42, Japanese quail, Mammalia, chick, human, mouse, rat | 26 | 0.683 | 0.869 |
| Mammalia, dog, golden (Syrian) hamster, human, mouse, rabbit, rat | 12 | 0.674 | 0.911 |

| | | | |
|---|---|---|---|
| Mammalia, human, mouse, rat | 13 | 0.688 | 0.899 |
| Mammalia, clawed frog, human, mouse | 32 | 0.699 | 0.997 |
| Japanese pufferfish, Mammalia, chick, guinea pig, human, mouse, rat, zebra fish | 14 | 0.605 | 0.752 |
| | 8 | 0.825 | 0.845 |
| human, mouse | 15 | 0.852 | 0.903 |
| Mammalia, human, mouse, rat | 12 | 0.916 | 0.906 |
| Mammalia, chick, clawed frog, human, mouse, rat | 25 | 0.664 | 0.94 |
| Mammalia, canary, chick, chimpanzee, clawed frog, common marmoset, dog, domestic pig, gibbon ape, goldfish, human, monkey, mouse, rainbow trout, rat, sheep, wild cat, woodchuck, zebra fish | 19 | 0.7 | 0.988 |
| Mammalia, cat, chick, clawed frog, domestic pig, golden (Syrian) hamster, hamster, human, monkey, mouse, rabbit, rat, sheep | 8 | 0.595 | 0.62 |

| | | | |
|---|---|---|---|
| Mammalia, chick, human, mouse, rabbit, rat | 79 | 0.659 | 0.856 |
| Mammalia, human, mouse, rat | 31 | 0.797 | 0.976 |
| Mammalia, cattle, chick, horse, human, monkey, mouse, rabbit, rat | 75 | 0.658 | 0.839 |
| human, mouse, rat | 164 | 0.709 | 0.941 |
| Chinese hamster (Gray hamster), Mammalia, cattle, chick, clawed frog, golden (Syrian) hamster, hamster, human, long-tailed hamster, monkey, mouse, rat, sheep | 41 | 0.812 | 0.961 |
| Mammalia, chick, golden (Syrian) hamster, human, mouse | 23 | 0.716 | 0.883 |
| Mammalia, chick, clawed frog, human, mouse, rat | 13 | 0.7 | 0.957 |
| Mammalia, human, mouse, rat | 19 | 0.779 | 0.843 |
| Chinese hamster (Gray hamster), Mammalia, human, mouse, rat | 14 | 0.682 | 0.85 |
| Chinese hamster (Gray hamster), Mammalia, chick, clawed frog, gibbon ape, golden (Syrian) hamster, human, monkey, mouse, rat | 44 | 0.627 | 0.768 |
| Chinese hamster (Gray hamster), Mammalia, chick, clawed frog, gibbon ape, golden (Syrian) hamster, human, monkey, mouse, rat | 51 | 0.769 | 0.93 |

| | | | |
|---|---|---|---|
| Chinese hamster (Gray hamster), Mammalia, chick, clawed frog, gibbon ape, golden (Syrian) hamster, human, monkey, mouse, rat | 18 | 0.689 | 0.746 |
| Chinese hamster (Gray hamster), Mammalia, chick, clawed frog, gibbon ape, golden (Syrian) hamster, human, monkey, mouse, rat | 54 | 0.753 | 0.906 |
| Mammalia, human, monkey, mouse, rat | 0 | 0.744 | 0.828 |
| Mammalia, human, monkey, mouse, rat | 0 | 0.535 | 0.786 |
| Chinese hamster (Gray hamster), Mammalia, cat, chick, clawed frog, gibbon ape, golden (Syrian) hamster, human, monkey, mouse, rat | 55 | 0.645 | 0.845 |
| Mammalia, mouse | 38 | 0.717 | 0.962 |
| Mammalia, cattle, human, mouse, rat | 10 | 0.679 | 0.877 |
| human, mouse, rat | 16 | 0.707 | 0.821 |
| Beechey ground squirrel, Chinese hamster (Gray hamster), Mammalia, cat, cattle, chick, clawed frog, dog, golden (Syrian) hamster, horse, human, macaque, medaka, monkey, mouse, pig, rabbit, rainbow trout, rat, sheep, zebra fish | 20 | 0.755 | 0.908 |
| Mammalia, chick, human, mouse | 32 | 0.79 | 0.837 |
| Mammalia, chick, human, mouse, rat | 26 | 0.515 | 0.697 |
| human, mouse, rat | 17 | 0.595 | 0.88 |
| human, mouse, rat | 24 | 0.766 | 0.81 |
| Japanese pufferfish, Mammalia, chick, human, mouse, zebra fish | 7 | 0.617 | 0.717 |

| | | | |
|---|---|---|---|
| Japanese pufferfish, Mammalia, chick, human, mouse, zebra fish | 5 | 0.54 | 0.655 |
| Japanese quail, Mammalia, chick, clawed frog, hamster, human, mouse, quail, rabbit, rat, zebra fish | 22 | 0.493 | 0.685 |
| Japanese pufferfish, Mammalia, human, mouse, rat, zebra fish | 35 | 0.697 | 0.81 |
| Mammalia, human, mouse, rat | 40 | 0.605 | 0.83 |
| Mammalia, human, mouse, rat | 9 | 0.617 | 0.841 |
| Mammalia, human, mouse, rat | 42 | 0.735 | 0.745 |
| Mammalia, chick, goldfish, human, mouse, rat | 17 | 0.777 | 0.851 |
| human, mouse, rat | 17 | 0.512 | 0.702 |
| human, mouse, rat | 7 | 0.511 | 0.709 |
| human, mouse, rat | 16 | 0.634 | 0.828 |
| Chinese hamster (Gray hamster), Mammalia, Vertebrata, cattle, clawed frog, human, monkey, mouse, rabbit, rat | 18 | 0.662 | 0.814 |
| Mammalia, clawed frog, human, mouse, rat | 7 | 0.637 | 0.749 |
| Mammalia, clawed frog, human, monkey, mouse, quail, rat | 19 | 0.644 | 0.711 |
| Chinese hamster (Gray hamster), Mammalia, Vertebrata, cattle, clawed frog, human, monkey, mouse, rabbit, rat | 72 | 0.605 | 0.697 |
| Chinese hamster (Gray hamster), Mammalia, Vertebrata, cattle, clawed frog, human, monkey, mouse, rabbit, rat | 28 | 0.433 | 0.596 |
| EBV | 35 | 0.754 | 0.91 |

| | | | |
|---|---|---|---|
| Mammalia, human, monkey, mouse, rabbit, rat | 11 | 0.7 | 0.883 |
| Mammalia, human, mouse, rat | 10 | 0.814 | 0.951 |
| Mammalia, human, mouse | 12 | 0.806 | 0.961 |
| Mammalia, human, mouse, rat | 44 | 0.907 | 0.957 |
| Mammalia, human, mouse, rat | 27 | 0.639 | 0.841 |
| human, mouse, rabbit | 59 | 0.911 | 0.968 |
| Mammalia, cattle, chick, domestic pig, frog, horse, human, medaka, mouse, rat, turtle, zebra fish | 8 | 0.7 | 0.927 |
| Mammalia, cattle, chick, dog, domestic pig, human, mouse, rat, zebra fish | 9 | 0.629 | 0.804 |
| Mammalia, chick, human, mouse, rat | 73 | 0.677 | 0.808 |
| Chinese hamster (Gray hamster), Mammalia, cattle, chick, clawed frog, dog, domestic pig, golden (Syrian) hamster, hamster, human, mink, monkey, mouse, rabbit, rat, sheep | 108 | 0.734 | 0.881 |
| Mammalia, cattle, chick, dog, golden (Syrian) hamster, human, monkey, mouse, rabbit, rat, sheep | 6 | 0.662 | 0.777 |
| human, mouse, rat | 30 | 0.761 | 0.948 |
| Chinese hamster (Gray hamster), Mammalia, cattle, chick, domestic pig, human, monkey, mouse, rabbit, rat | 30 | 0.666 | 0.707 |
| Chinese hamster (Gray hamster), Mammalia, cattle, chick, domestic pig, human, mink, monkey, mouse, pig, rabbit, rat | 11 | 0.586 | 0.841 |
| Chinese hamster (Gray hamster), Mammalia, cattle, chick, domestic pig, human, mink, monkey, mouse, pig, rabbit, rat | 68 | 0.793 | 0.879 |

| | | | |
|---|---|---|---|
| Mammalia, cat, cattle, chick, clawed frog, dog, human, mouse, rat | 0 | 0.787 | 0.874 |
| Mammalia, cat, cattle, chick, clawed frog, dog, human, mouse, rat | 21 | 0.688 | 0.915 |
| Mammalia, clawed frog, human, mouse, rat | 10 | 0.793 | 0.888 |
| cattle, human, monkey, mouse, rat | 55 | 0.384 | 0.745 |
| Mammalia, cattle, dog, human, long-tailed hamster, monkey, mouse, rabbit, rat, sheep | 30 | 0.738 | 0.872 |
| human | 12 | 0.713 | 0.817 |
| Chinese hamster (Gray hamster), Mammalia, chick, golden (Syrian) hamster, human, mouse, rat | 44 | 0.669 | 0.85 |
| HTLV-I, Mammalia, cattle, domestic pig, hamster, human, mink, monkey, mouse, rabbit, rat, sheep | 17 | 0.733 | 0.771 |
| HTLV-I, Mammalia, cattle, domestic pig, hamster, human, mink, monkey, mouse, rabbit, rat, sheep | 5 | 0.5 | 0.643 |
| Mammalia, chick, clawed frog, human, mouse, rat | 20 | 0.764 | 0.925 |
| Mammalia, human, mouse, rat | 41 | 0.615 | 0.885 |
| Mammalia, human, mouse, rat | 31 | 0.701 | 0.882 |
| Mammalia, clawed frog, dog, human, mouse, rat | 6 | 0.688 | 0.962 |
| human | 5 | 0.5 | 0.752 |
| Mammalia, cattle, human, mouse, rat | 28 | 0.728 | 0.946 |
| human, mouse, rat | 6 | 0.705 | 0.815 |

| | | | |
|---|---|---|---|
| Mammalia, cattle, human, mouse, pig, rat, sheep | 6 | 0.673 | 0.973 |
| chick, domestic pig, human, mouse, rat | 12 | 0.657 | 0.824 |
| ASV 17 | 24 | 0.7 | 0.895 |
| AMV | 29 | 0.669 | 0.835 |
| Mammalia, human, mouse, rat | 23 | 0.967 | 0.982 |
| clawed frog | 6 | 0.609 | 0.783 |
| clawed frog | 8 | 0.634 | 0.833 |
| Mammalia, cattle, chick, human, mouse, rat | 18 | 0.721 | 0.916 |
| Mammalia, cattle, chick, human, mouse, rat | 23 | 0.688 | 0.843 |
| human, mouse, rat | 17 | 0.718 | 0.759 |
| human, mouse | 35 | 0.648 | 0.846 |
| human, mouse | 35 | 0.749 | 0.885 |
